# Kinase Plasticity with Vandetanib Treatment Enhances Sensitivity to Tamoxifen in Estrogen Receptor Positive Breast Cancer

**DOI:** 10.1101/2024.12.19.629395

**Authors:** Austin A. Whitman, Rasha T. Kakati, Santiago Haase, Christine HNC Thai, Attila T. Szenasi, Elizabeth C. Brunk, Denis O. Okumu, Michael P. East, Sarahi G. Molina, Adriana S. Beltran, Jesse R. Raab, Lisa A Carey, Charles M. Perou, Gary L. Johnson, Philip M. Spanheimer

**Author notes:** AAW, RTK, and SH contributed equally to this manuscript. Correspondence should be addressed to: Philip M. Spanheimer MD, Lineberger Comprehensive Cancer Center, University of North Carolina at Chapel Hill, 170 Manning Drive, Suite 1149, Chapel Hill, NC 27599-7213 USA. Disclosure of conflict of interests: C.M.P is an equity stockholder and consultant of BioClassifier LLC; C.M.P is also listed as an inventor on patent applications for the Breast PAM50 Subtyping assay. The remaining authors declare no potential conflicts of interest.

## Abstract

Resistance to endocrine therapy (ET) is common in estrogen receptor-positive (ER+) breast cancer. Multiple studies have demonstrated that upregulation of MAPK signaling pathways contributes to ET resistance. Herein we show that vandetanib treatment suppresses MAPK signaling and enhances sensitivity to ET across ET-sensitive and ET-resistant ER+ cell lines and patient derived organoids. Vandetanib treatment reprograms transcription toward a less proliferative, more estrogen responsive, Luminal-A like state by enriching ERα chromatin binding at canonical estrogen response elements. Multiplexed kinase inhibitor beads-mass spectrometry (MIB/MS) revealed kinase network reprogramming, including upregulation of PI3K and HER2 activity, as shared adaptive resistance mechanisms to vandetanib treatment. Co-treatment with the HER2 inhibitor lapatinib, further enhanced sensitivity to vandetanib. Using an operating room-to-laboratory short-term *ex-vivo* assay coupled to single-cell RNA sequencing, we demonstrate conserved gene expression changes in primary tumor cells, including increased HER2 activity signatures, following vandetanib treatment. Vandetanib sensitivity signatures were generated from cell line and primary human tumor cells which correlate with vandetanib sensitivity in ER+ patient-derived organoid and xenograft models. Future clinical trials of vandetanib in ER+ breast cancer should include rationally designed co-treatments based on adaptive resistance pathways, including HER2, and evaluate response signatures as biomarkers predicting patients most likely to benefit.

**SIGNIFICANCE:** Vandetanib enhances sensitivity to tamoxifen in ER+ breast cancer by reprograming ERα gene regulation and kinase signaling networks which define gene expression signatures associated with response and identify targetable adaptive resistance pathways.

## INTRODUCTION

Approximately 75% of breast cancers are estrogen receptor-positive (ER+) and fall within the luminal subtypes (1-3). Most patients with ER+ breast cancer receive endocrine therapy (ET) targeting ER. However, resistance to ET occurs in about 1/3 of early cancers and eventually develops in all metastatic ER+ cancers(4-6). Targeted inhibitors to PI3K, mTOR, and CDK4/6 have shown efficacy in patients with ET-resistant disease (7-10), but responses are often short lived, and many patients remain in need of durable, effective therapy.

Activation of the ERK/MAPK pathway is a well-established mechanism by which ER+ breast tumors resist ET (11-17). Gain-of-function mutations in ERK/MAPK nodes or upstream receptor tyrosine kinase (RTK) are enriched in ET-resistant tumors, although these changes occur in only ∼13% of cases (11). ERK/MAPK can be activated by upstream recpetors, including the RET receptor tyrosine kinase, which is expressed in ∼50% of ER+ breast tumors and is more highly expressed in ET-resistant tumors (18-21). Co-expression of RET and its activating ligand, glial derived neurotropic factor (GDNF), is associated with transciptional signatures of ERK/MAPK activation and with worse outcomes in human ER+ breast tumors (22). Further, RET is overexpressed with ESR1 (encoding ER) activating mutations, and inhibiting RET in ESR1 mutant models suppresses tumor growth (23).

Our prior work showed that tyrosine kinase inhibitors (TKIs) that inhibit RET, such as vandetanib and sunitinib, reduce proliferation and enhance ET sensitivity in ER+ breast cancer cell lines (24, 25). In RET-positive primary breast tumors, *in vitro* TKI treatment reduced ERK activity (26). These findings support a role for TKIs to inhibit ERK/MAPK activation to improve response to ET.

Despite encouraging preclinical data, clinical trials with vandetanib have had disappointing results. However, these trials failed to account for potentially predictive biomarkers such as ER and RET or evaluated vandetanib only as a monotherapy. For example, a phase II trial of single-agent vandetanib in 46 patients with metastatic breast cancer did not demonstrate a benefit (27), but did not report receptor status, limiting the interpretation of whether this trial included patients that mechanistically would be predicted to respond. Our work, including the findings presented herein, suggests that vandetanib may provide a benefit for select patients. Recent TKI trials enrolling only ER+ patients have shown promise (28, 29), renewing interest in TKI and highlighted the need for biomarkers to identify patients most likely to respond to TKI therapy. To develop biomarkers of response and to elucidate bypass resistance mechanisms, we sought to investigate how vandetanib remodels gene expression and kinase signaling pathways in ER+/HER2- breast cancer.

## MATERIALS AND METHODS

### Cell Lines

We obtained ER+ breast cancer cell lines parental MCF7 (RRID: CVCL_0031), tamoxifen-resistant MCF7 (MCF7-TAM) (RRID:CVCL_1D43), exemestane-resistant MCF7 (MCF7-EXE) (RRID:CVCL_M436), parental T47D (RRID:CVCL_0195), and tamoxifen-resistant T47D (T47D-TAM) (RRID:CVCL_1D36), from the American Type Culture Collection (ATCC). Cell lines were cultured using Dulbecco’s Modified Eagle Medium (DMEM) (Gibco) with 10% fetal bovine serum (Gibco) and 1% penicillin-streptomycin (Life Technologies) at 37°C and 5% CO2. MCF7-TAM and T47D-TAM were maintained by chronic treatment with 1 μM of 4OH-Tam and MCF7-EXE with 0.1 μM of exemestane.

### Chemicals and Treatment

Vandetanib (SelleckChem) and z-4-Hydroxytamoxifen (4OH-Tam) (SelleckChem) were dissolved in dimethyl sulfoxide (DMSO). Pralsetinib was obtained from Chemietek, and lapatinib from Medchem Express LLC. Treatments were performed for 24 hours unless otherwise specified.

### Cell Viability Assays

Cells were plated in opaque-walled 96-well plates at a density of 4,000 cells per well in technical triplicates and allowed to adhere overnight before the addition of the specified inhibitors. Viability was assessed using the CellTiter-Glo (CTG) Luminescent Cell Viability Assay (Promega). Luminescence was measured using a Synergy 2 microplate reader. Samples were averaged across biological triplicates. Statistical analysis and plotting were performed using GraphPad Prism 9.

### Western Blots

Cells were lysed in Pierce RIPA buffer containing Halt protease inhibitor cocktail and PhosStop phosphatase inhibitor (Thermo Scientific). Proteins were resolved by SDS-polyacrylamide gel electrophoresis, transferred to PVDF membrane, and blocked with 10% BSA in PBS, then incubated overnight with the indicated primary antibody in blocking buffer. Membranes were washed three times in Tris-buffered saline containing 0.1% tween-20 then incubated for 1 hour with HRP-conjugated secondary antibodies. Blots were developed using a ChemiDoc-touch digital imager (Bio-Rad) and band intensity quantified using ImageJ (https://imagej.net/ij/index.html). The antibodies included: p44/42 MAPK (ERK1/2) (Cell Signaling Technology Cat# 9107, RRID:AB_10695739), Phospho-p44/42 MAPK (Erk1*/*2) (Cell Signaling Technology Cat# 75165, RRID:AB_2728835), RET (Abcam Cat# 3454-1, RRID:AB_10639232), and β-Actin (Santa Cruz Biotechnology Cat# sc-47778, RRID:AB_626632).

### Patient Derived Organoid Derivation

Samples were obtained from patients with breast cancer who were undergoing surgical resection. This was performed under an IRB-approved protocol with written informed consent to generate new patient-derived breast cancer models. Patient-derived organoids (PDOs) were established from human tumor tissue samples as previously described (30). Briefly, tumor tissues were finely minced and digested enzymatically with a collagenase–hyaluronidase mixture (STEMCELL Technologies, Cat. No. 07912) for 2–5 h at 37 °C to release viable cell clusters. Red blood cells were removed using ACK Lysing Buffer (Gibco, Cat. No. A10492-01). The resulting cell clusters were resuspended in growth factor–reduced Matrigel (Corning, Cat. No. 356231) and plated as 30 µL domes in prewarmed 6-well plates. After Matrigel polymerization, domes were overlaid with breast cancer organoid (BCO) medium composed of Advanced DMEM/F-12 (Gibco, Cat. No. 12634010) supplemented with 1X GlutaMAX (Gibco, Cat. No. 35050061), 10 mM HEPES (Gibco, Cat. No. 15630080), 1X Penicillin–Streptomycin (Gibco, Cat. No. 15140122), 50 µg/mL Primocin (InvivoGen, Cat. No. ant-pm-05), 1X B27 Supplement (Gibco, Cat. No. 17504001), 5 mM Nicotinamide (Thermo Fisher Scientific, Cat. No. 128271000), 1.25 mM N-acetyl-L-cysteine (MilliporeSigma, Cat. No. A9165), 250 ng/mL R-spondin-3 (PeproTech, Cat. No. 120-44-100UG), 5 nM Heregulin β1 (PeproTech, Cat. No. 100-03-100UG), 100 ng/mL Noggin (PeproTech, Cat. No. 120-10C-100UG), 20 ng/mL FGF-10 (PeproTech, Cat. No. 100-26-100UG), 5 ng/mL FGF-7/KGF (PeproTech, Cat. No. 100-19-100UG), 5 ng/mL EGF (PeproTech, Cat. No. AF-100-15-100UG), 500 nM A-83-01 (Selleck Chemicals, Cat. No. S7692), and 500 nM SB202190 (Selleck Chemicals, Cat. No. S1077). Y-27632 (Selleck Chemicals, Cat. No. S1049; 5 µM) was included for the first three days of culture to enhance cell survival. Culture medium was refreshed every 3–4 days.

PDOs were passaged every 2–4 weeks. Matrigel domes were collected, mechanically disrupted, and incubated with TrypLE™ Express (Gibco, Cat. No. 12604013) for 7–10 min at 37 °C. The enzyme was neutralized with soybean trypsin inhibitor (Gibco, Cat. No. 17075029). Cells were then washed with Advanced DMEM/F-12, resuspended in cold Matrigel, and replated to generate new organoids. After each passage, dissociated cells were plated for immunofluorescence.

### Immunofluorescence Staining

PDOs were dissociated, and single cells were seeded in 96-well plates for 5–24 h. Cells were fixed with 4% paraformaldehyde (Electron Microscopy Sciences, Cat. No. 15710) in PBS for 15 min at room temperature, then washed three times with PBS. Samples were permeabilized with 0.3% Triton™ X-100 (MilliporeSigma, Cat. No. T8787) in PBS for 15 min. Non-specific binding was blocked with 5% bovine serum albumin (BSA; MilliporeSigma, Cat. No. A9647) in PBS for 1–2 h at room temperature. Primary antibodies were diluted in 1% BSA/0.1% Triton X-100/PBS and incubated overnight at 4 °C: anti-Estrogen Receptor α (ERα; Abcam, Cat. No. ab32063, RRID: AB_732249) and anti-ERα, clone SP1 (Thermo Fisher Scientific, Cat. No. RM-9101-S, RRID: AB_720919), each at 1:200. The following day, cells were washed three times (5 min each) in 1% BSA/0.1% Triton X-100/PBS and incubated for 1 h at room temperature in the dark with Alexa Fluor™–conjugated secondary antibodies (Thermo Fisher Scientific, Cat. Nos. A-21236 and A-21244, RRIDs: AB_2535805 and AB_2535812, respectively) at 1:500 dilution in blocking buffer. Nuclei were counterstained with DAPI (1:5000 from a 10 µg/mL stock; Thermo Fisher Scientific, Cat. No. D1306) in PBS. After three final PBS washes, cells/organoids were maintained in PBS or mounting medium and imaged using an EVOS™ Auto fluorescence microscope. Secondary-only controls were included to verify signal specificity.

### Drug Treatment and Viability Assay

For drug treatment, organoids were dissociated into single cells by mechanical disruption of Matrigel domes, followed by incubation with TrypLE™ Express (Gibco, Cat. No. 12604013) for 8 min at 37 °C. The enzyme was neutralized, and cells were pelleted by centrifugation at 300 × g for 5 min, then resuspended in ice-cold growth factor–reduced Matrigel (Corning, Cat. No. 356231). For each well, 5,000 viable cells were seeded in 10 µL Matrigel and plated in the center of a white-walled 384-well tissue culture plate to form a dome. Plates were incubated at 37 °C for 15–30 min to allow Matrigel polymerization, then 40 µL of prewarmed BCO medium was gently added to each well. Drug treatments were initiated at 72 h after seeding. Organoids were exposed to varying concentrations of tamoxifen or vandetanib, either alone or in combination, with vehicle control containing dimethyl sulfoxide (DMSO; Sigma-Aldrich, Cat. No. D2650). Each treatment was performed in technical and biologic triplicate. Cell viability was assessed 72 h post-treatment using the CellTiter-Glo 3D Cell Viability Assay (Promega, Cat. No. G9682, RRID: SCR_006724). Plates were shaken on an orbital shaker for 30 min at room temperature. Luminescence was measured using a microplate reader (BMG Labtech CLARIOstar), and background signal from Matrigel-only wells was subtracted.

### RET CRISPR Knockout

Two sgRNAs targeting exon 3 of RET (5’-ACTGGCACTCGCCCTCACGA-3’ and 5’-GGGGTCGGTTCTCCCGAATG-3’), according to UCSC Genome Browser, were cloned into the pSpCas9(BB)-2A-Puro(PX459) V2.0 plasmid (Addgene) and verified by Sanger sequencing. MCF-7 cells were cultured in antibiotic-free media and grown to 70-90% confluency. Cells were then transfected with sgRNA plasmids using FuGENE 4K Transfection Reagent according the manufacturer’s recommendations. After 72 hours, cells were passaged onto 15-cm plates at a low density and selected with 1 μg/mL puromycin (Gibco) for 3 days. Cell colonies were then isolated and genomic DNA was purified using the Quick-DNA Miniprep Kit (Zymo Research). The targeted region was then amplified by PCR and sequenced confirming frameshift mutations resulting in the introduction of premature stop codons in both sgRNA-targeted alleles.

### Synergy Analysis

Dose–response curves for tamoxifen, vandetanib, and vandetanib plus 0.5 µM tamoxifen were fitted using a four□parameter logistic model (4PL). From each curve, we extracted the drug concentration required to reach 50% inhibition.

Combination Index (CI50)

CI50 = (D1 / Dx1) + (D2 / Dx2)

D1 = vandetanib concentration in the combination

Dx1 = vandetanib concentration alone

D2 = fixed tamoxifen dose (0.5 µM)

Dx2 = tamoxifen concentration alone at 50% inhibition

CI < 1 indicates synergy, CI ≈ 1 additivity, CI > 1 antagonism.

Sensitization Ratio (SR50)

SR50 = Dx1 / D1

SR > 1 indicates increased vandetanib potency in the presence of tamoxifen.

### mRNA Sequencing and Data Analysis

Cells were treated with 10 µM vandetanib or vehicle for 24 hours in biological triplicates and total RNA was isolated (Qiagen). mRNA sequencing libraries were made using TruSeq RNA Library Prep Kit (Illumina) and sequenced using the Illumina NextSeq2000. Plots and statistical analyses were performed using RStudio 2022.07.1 Build 554. Differential gene expression (DGE) was determined using the DESeq2 R package using raw mRNA-seq counts from STAR/Salmon. Genes with less than 10 counts across all samples were omitted before analysis and counts were filtered and normalized to the upper quartile using NOISeq. Significant differentially expressed genes were defined by using a Log2FC cutoff of +/- 0.5 and a false discovery rate (FDR) < 0.05. Enrichment of gene sets was investigated through hypergeometric analysis of an unranked list of upregulated or downregulated elements using the enricher function of the clusterProfiler package. METABRIC and SCAN-B databases were obtained for survival analysis and filtered for ER+/HER2- patients treated with ET. A median-centroid approach was implemented to assign a gene set score per patient. Kaplan-Meier (KM) plots were then computed using the survminer and survival packages.

### Cleavage Under Targets and Release Using Nuclease (CUT&RUN) and Analysis

Cut and Run was performed with the EpiCypher Cutana CUT&RUN Kit Version 3 according to the manufacturer’s manual v3.5. Cells were treated with 10 µM vandetanib or vehicle for 24 h, and cell counts were quantified using the Countess. 500,000 viable cells were bound to ConA beads per reaction, and 0.5 ug (or manufacturer’s recommendation) of antibody was added to each sample overnight. pAG-MNase was then used to cleave DNA, which was extracted and used to generate libraries following the EpiCypher Cutana CUT&RUN Library Prep Kit. Libraries were sequenced on the NextSeq2000 (Illumina) platforms with paired-end sequencing and dual indexing, according to the manufacturer’s protocol.

Cut and Run sequencing results were processed through the Nextflow DSL2 implementation of the Raab Lab CUT&RUN processing pipeline (https://github.com/raab-lab/cut-n-run). The pipeline produced peak filtered calls and BAM coverage files, which were used to run differential peak binding between vandetanib versus vehicle treated samples for ERα, FOXA1 and H3K4me3 separately. For this, a union peak set was generated from the vehicle and vandetanib samples (with 2 replicates each) using bedtools merge function (31).

Peak count with IgG subtraction was performed on the union peakset by subtracting the corresponding IgG sequencing controls for Vehicle and Vandetanib-treated samples. Counts were library-size normalized with DiffBind dba.normalize(), and differential peak analysis was performed with DiffBind dba.analyze() with EdgeR (32). Gene annotations were added to the differential peak analysis using ChIPseeker annotatePeak()(33, 34). To analyze peak gene signature enrichment, the binding fold changes for all the peaks associated with a signature were computed (vandetanib vs vehicle) and a Wilcoxon test was performed to determine significant differential deposition for the given signature.

To illustrate the global shifts in ERα, FOXA1 and H3K4me3 deposition, we extracted the Diffbind EdgeR top upregulated and downregulated differential peaks (vandetanib vs vehicle) and generated peak profile heatmaps from normalized coverage plots using deepTools computeMatrix and plotHeatmap functions (35). For significant genes and peaks, coverages plots were produced using library size normalized coverage plots with Gviz GeneRegionTrack (36). To illustrate peak gene distribution, we used the CnRAP (Cut & Run Analysis Pipeline) step 4 script, which generates peak gene distribution images from peak annotations using a modified CUT&RUN analysis pipeline to identify transcription factor binding sites in human cell lines (37). The CUT&RUN quality assessment through peak profile heatmaps, transcriptional start site distribution and peak gene distribution are in Fig 5E-G.

### MIB/MS and Data Analysis

Cells were treated with vandetanib at its IC90 for 48 h. Cells were harvested by scraping and cell pellets obtained for proteomic analysis using the MIB/MS technology (38, 39). Scaled label free quantification (LFQ) values were obtained and a correlation plot of samples was computed to assess homogeneity of triplicates and heterogeneity of treatment conditions for quality control using ggplot2. Differential protein expression (DPE) was analyzed using the limma R package using scaled LFQ values. Significant differentially expressed proteins were defined using a Log2FC (log2 fold change) cutoff of 0.5 for upregulation and -0.5 for downregulation, along with a p-value < 0.05.

### Primary ER+/HER2- Human Tumor Cells

A primary tumor sample was obtained from a patient with ER+/HER2- invasive ductal carcinoma. The tumor was processed fresh as previously described (40). In brief, tumor tissue was transported to the laboratory immediately after surgical resection, dissociated into a single cell suspension, and treated with vandetanib (10 µM) or control media for 12 hours. After treatment, single cell RNA sequencing libraries were made using the 10x Genomics Chromium Next GEM Single Cell 3’ v3.1 and sequenced on the NextSeq2000 (Illumina) with pair-end sequencing.

### Single Cell RNA Sequencing Analysis

Reads were aligned and counted using Cellranger count function with default parameters. The counts and features matrixes were used to generate Seurat objects. The Seurat objects corresponding to each treatment group were merged with Seurat’s function merge (), and low quality cells were removed. Doublets were removed using scDblFinder (41) and cells were filtered: nFeature_RNA > 200 & nFeature_RNA < 6000 & percent.mt < 10. Cell types were annotated on the combined Seurat object using SingleR using the cell type atlas HumanPrimaryCellAtlasData () accessed with the package celldex. InferCNV was used to predict copy number alterations, defining the immune cells ("T_cells","Macrophage","DC","Monocyte","NK_cell") as reference.

Differential analysis of gene expression was performed on the tumor cells to compare vandetanib-treated versus non-treated cells using findmarkers (). GSEA was then performed using the list of differentially expressed genes, ranking the genes according to log2(FC)/FDR, and the ranked gene lists were used in the GSEA software (42). Gene signatures were scored using AddModuleScore_UCell() from the UCell package.

We used the NMF R package to build a non-negative matrix factorization (NMF) model of the tumor cells using a list of genes differentially expressed upon vandetanib treatment in MCF7 cells from this study. The Seurat object was subset to contain only the above-mentioned genes, and the NMF model was fitted to 8 factors. Heatmaps were generated for each treatment group to assess enrichment of cells to different NMF factors. Each cell was assigned to a unique NMF according to the NMF with maximum coefficient value for that cell, and the percentage of cells belonging to each NMF factor was calculated for each treatment group. DE analysis was performed comparing cells in certain NMF factors.

### Patient Derived Xenografts

Two ER+ patient-derived xenografts (PDX) were obtained as a generous gift from Dr. Alana Welm (43). PDXs were orthotopically injected into athymic mice that had been implanted with estrogen releasing pellets (Innovative Research of America, NE-121). When tumors were 0.5 cm in greatest dimension, five mice were randomized to each of the following treatments: control gavage, tamoxifen (50 mg/kg daily), or vandetanib (25 mg/kg daily). Tumor size was measured three times per week using calipers. After 6 weeks, mice were sacrificed and tumors were collected for mRNA sequencing.

### Statistics

All experiments were performed with at least 3 biological or technical replicates. Small sample number comparisons were analyzed with an unpaired *t* test with Welch’s correction for non-equal SD. Linear regressions and nonlinear regression curves were compared by statistical tests on the parameters of the curves using the extra sum of squares *F* test. Experiments with large sample numbers were analyzed by Wilcoxon test.

### Data Availability

Single cell and bulk RNA sequencing data was made available at the Sequence Read Archive (SRA) under BioProject PRJNA1177387. Mass spectrometry protein sequencing results are included in this manuscript as Supplementary Table 1.

### Code Availability

All original code has been deposited on the Github, and it is publicly available at the Github repository at https://github.com/santiagohaase/Spanheimer-lab-Kinase-Plasticity-in-Response-to-Vandetanib-Enhances-Sensitivity-to-Tamoxifen

## RESULTS

### Vandetanib Inhibits ERK Activation and Viability in ER+ Breast Cancer Cells

Vandetanib inhibits receptor tyrosine kinases (RTK) including RET and EGFR which are upstream activators of ERK/MAPK (Fig. 1A). We first investigated if vandetanib inhibited the ERK/MAPK pathway using MCF7 and two ET-resistant derivatives, MCF7-TAM (tamoxifen) and MCF7-EXE (exemestane), which had been cultured long-term under drug treatment to develop resistance. Vandetanib treatment reduced both the cellular protein levels of ERK and levels of activating phosphorylation of ERK (Fig. 1B). To account for changes in total ERK, we calculated the ratio of pERK/ERK signal intensity, which was reduced in all 3 cell lines with vandetanib treatment. Dose-response curves demonstrated dose IC50 values for vandetanib of 8.5 μM in MCF7, 10.4 μM in MCF7-TAM, and 15.9 μM in MCF7-EXE (Fig. 1C). Similar vandetanib sensitivities were seen in T47D and T47D-TAM (Fig. S1).

**Figure 1.**
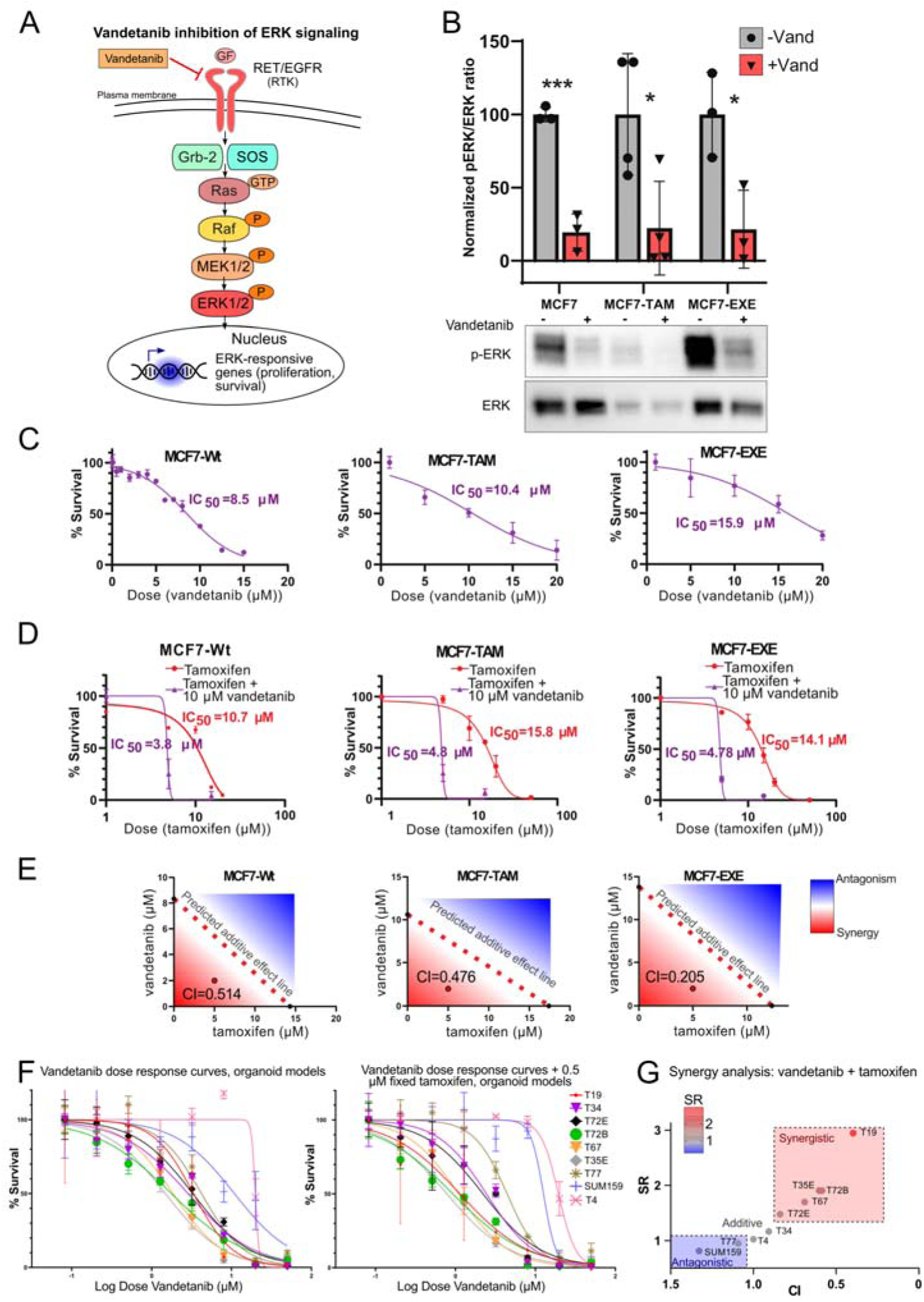
Vandetanib reduces ERK activity and is synergistic with Tamoxifen in endocrine sensitive and resistant breast cancer cells and Patient Derived Organoids. (A) Schematic of vandetanib target kinases and downstream RAS-RAF-ERK signaling cascade. (B) Treatment of MCF-7 and tamoxifen resistant MCF7 (MCF7-TAM) and exemestane resistant MCF7 (MCF7-EXE) reduces pERK/ERK in all 3 cell lines. *p<0.05, ***p<0.01 by t-test with Welch correction (C) Vandetanib causes a dose dependent reduction in cell viability in MCF7, MCF7-TAM, and MCF7-EXE. (D) Treatment with Vandetanib results in a left shift of the tamoxifen sensitivity curve and reduction in IC50 in all 3 cell lines. (E) Isobologram plots showing predicted additive lines and regions of synergy (red) or antagonism (blue) for vandetanib–tamoxifen combinations demonstrate synergy in all 3 cells lines. CI = Combination Index (<1 indicates synergic interaction) (F) Patient derived breast cancer organoids show a range of vandetanib sensitiivty. (G) In total, 5 PDO showed synergy between vandetanib and tamoxifen, 2 had an additive effect, and 1 PDO and an organoid derived from the TNBC cell line SUM159 were antagonistic.

### Vandetanib Sensitizes ER+ Breast Cancer Cells to Tamoxifen

We next tested if co-treatment with vandetanib enhanced the antiproliferative effects of tamoxifen in ET-resistant cells. The IC50 of the active metabolite of tamoxifen, 4OH-Tam, was 10.7 μM in MCF7, 15.8 μM in MCF7-TAM, and 14.1 μM in MCF7-EXE (Fig. 1D) demonstrating decreased sensitivity to tamoxifen with chronic ET. Treatment with 10 μM vandetanib resulted in a left shift of the 4OH-Tam sensitivity curve in all 3 cell lines with the IC50 for 4OH-Tam of 3.8 μM in MCF7, 4.8 μM in MCF7-TAM, and 4.8 μM in MCF7-EXE. We produced an additive effect isobole line by connecting the individual tamoxifen and vandetanib concentrations that caused the survival percentage equal to the drug combination. The point indicating the combination of 2 μM of vantetanib with 5 μM of tamoxifen is below the additive isobole line, demonstrating a synergic effect for the MCF7, MCF7-TAM, and MCF7-EXE (Fig. 1E).

### Vandetanib Response in Breast Cancer Patient Derived Organoids

We derived a panel of primary breast tumor patient-derived organoids (PDOs) to investigate the impact of vandetanib across a diverse group of translationally relevant breast cancer models. Patient and tumor characteristics are listed with intrinsic subtype assigned from RNA-seq (Fig. S2A). Of the 6 ER+/HER2- PDOs, 2 were LumA subtype, 1 was LumB, 1 was HER2E, and 2 were basal-like subtype, recapitulating the molecular heterogeneity of ER+ disease. The TNBC derived PDO (T4) had a basal molecular subtype. PDOs displayed a range of sensitivity to tamoxifen (Fig. S2B), with ER-negative models, including the organoid derived from the TNBC cell line Sum149, having the lowest sensitivity. PDOs were further validated for expression of the estrogen receptor (ER), which displayed heterogeneity in ER expression within and across PDO, and the epithelial marker EPCAM (Fig. S2C). PDO displayed a range of sensitivity to vandetanib, with greater sensitivity in ER+ PDOs (Fig. 1F). To analyze the effect of the drug combination, we calculated the combination index (CI) and sensitization index (SR) at 50% of survival. To represent both the interaction (CI) and the magnitude of sensitization (SR), we plotted SR versus CI for each cell model at the 50% effect level. Low CI and high SR values indicate strong synergy. Synergy analysis demonstrated that 5 ER+ PDOs displayed synergy with dual treatment of vandetanib and tamoxifen, and this included 3 of 4 luminal intrinsic subtype (Fig. 1G). These results show a high vandetanib-tamoxifen synergy rate in PDOs that retain ER expression and luminal transcriptional profile.

### RET Regulates ERK Activity and Vandetanib Response in ER+ Breast Cancer Cells

Based on our prior work using a transient knockdown model showing a role for RET in mediating vandetanib response and driving poor outcomes and ET resistance in ER+ breast cancer (22), we explored the role of RET in vandetanib sensitivity. Transient RET antagonism with the selective RET inhibitor BLU667 resulted in reduced sensitivity to vandetanib in MCF7 cells in dose-dependent fashion (Fig. 2A). BLU667 also reduced vandetanib sensitivity in MCF7-TAM (Fig. S3A), but not in the low RET expressing MCF7-EXE (Fig. S3B).

**Figure 2.**
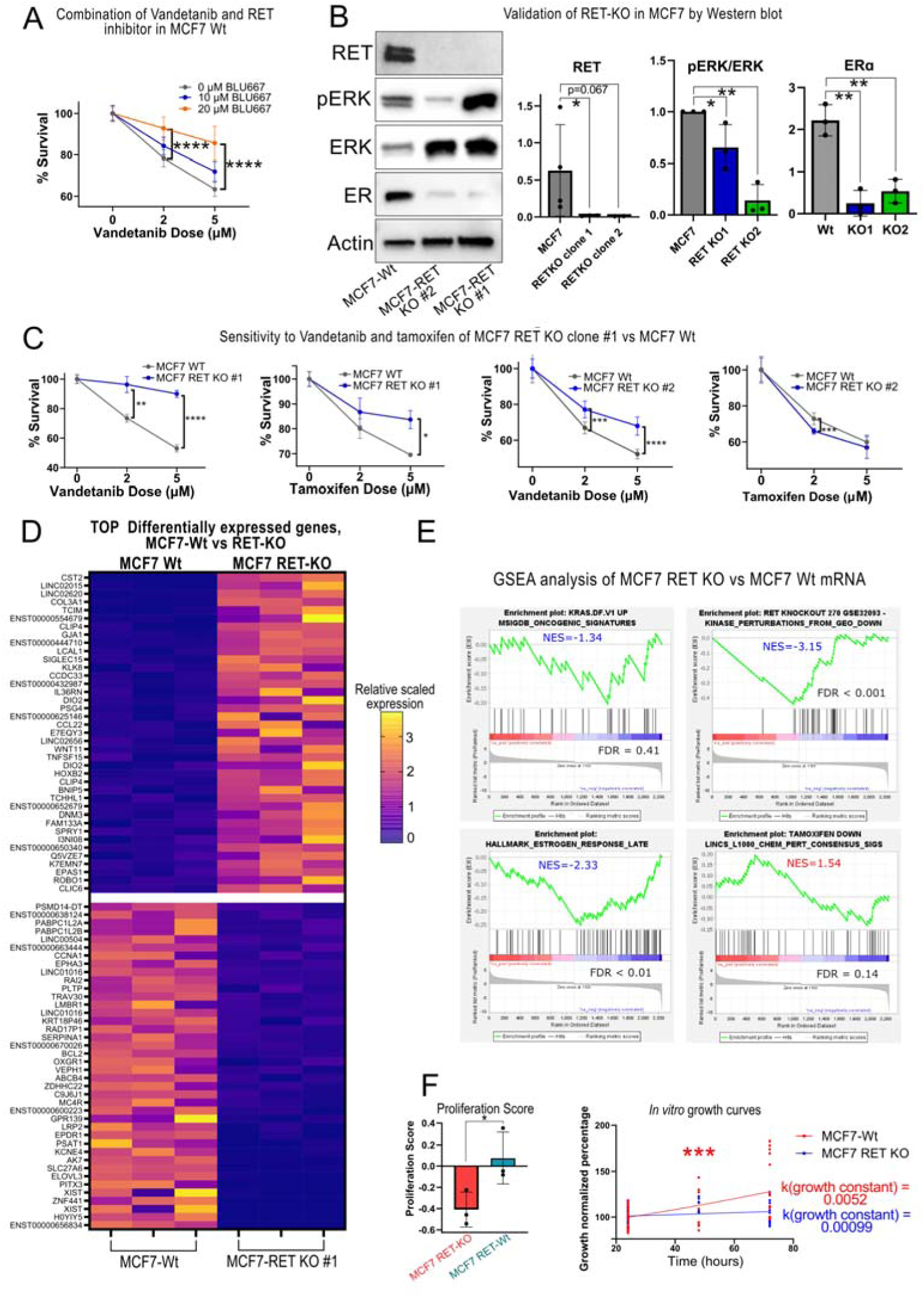
Vandetanib effects are partially mediated by the RET receptor. (A) Co-treatment with vandetanib and the selective RET inhibitor BLU667 reduced vandetanib sensitivity in MCF7(B) Western blot demonstrating CRISPR knockout of RET resulting in >90% reduction in RET protein levels. RET knockout resulted in reduced pERK and ER expression. (C) RET knockout cells have reduced sensitivity to vandetanib and tamoxifen. (D) Heatmap showing top up- and down-regulated genes in RET KO cells. (E) Gene Set Enrichment Analysis demonstrates reduction in hallmark RAS signaling and RET activity in RET KO cells. RET KO cells also show decreased expression of estrogen response genes consistent with reduced sensitivity to tamoxifen. (F) RET KO cells have lower proliferation score and reduced proliferation compared to WT. (. *p<0.05, **p<0.01, ***p<0.001, ****p<0.0001 by t-test with Welch correction for the bar graphs, and Wilcoxon test (A-C) and for statistical test on k (growth constant) parameter from lineal regression model (F).

To precisely elucidate the role of RET in ER+ breast cancer cells using a more robust experimental model than prior work transient siRNA knockdowns, we generated CRISPR knockouts of RET in MCF7 cells (RET KO#1 and RETKO#2). Western blot confirmed knockout of RET protein (Fig. 2B). Both knockouts had reduced pERK levels compared to wild type (WT), demonstrating that RET is a key regulator of this pathway. RET KO cells also had reduced ER protein levels. Further supporting a key role for RET as a mediator of vandetanib response, both KO lines were significantly less sensitive to vandetanib compared to WT (Fig. 2C). Based on the vandetanib synergy assay we hypothesized that RET KO would sensitize cells to tamoxifen, however, KO cells were less sensitive (Fig. 2C), likely because of reduced ER expression.

We further explored the molecular basis of this unexpected finding by comparing gene expression of RET KO vs WT cells using RNA sequencing. Differentially expressed gene analysis revealed many genes significantly regulated by RET expression (Fig. 2D) with RET KO. Hallmark gene sets for KRAS signaling and RET knockout were lower with RET KO (Fig. 2E), as expected. Consistent with reduced sensitivity to tamoxifen, estrogen response genes were reduced, indicating that RET KO cells have reduced ER activity. RET KO resulted in lower proliferation scores compared to WT (Fig. 2F) and RET KO cells had significantly reduced growth vs WT. Cumulatively, these findings support that RET promotes an ERK-driven proliferative phenotype and mediates vandetanib response, but stable knockout is associated with reduced expression of ER and estrogen response genes.

### Vandetanib Induces Low-Proliferative and Estrogen Responsive Gene Expression

To delineate the transcriptional basis for how vandetanib enhances tamoxifen sensitivity in ER+ breast cancer cells, we performed mRNA sequencing after treatment with vandetanib (10 μM) compared to vehicle. Heatmaps of DEGs are shown for MCF7 (Fig. 3A), MCF7-TAM (Fig. 3B), and MCF7-EXE (Fig. 3C). In MCF7, downregulated genes were enriched for MAPK signaling and proliferative signatures (Fig. 3D), consistent with loss of pERK observed in Fig. 1A. Upregulated genes are enriched for signatures of estrogen response and sensitivity to tamoxifen, consistent with observed sensitization to tamoxifen (Fig. 1D). Up-regulated and down-regulated gene sets in MCF7-TAM (Fig. 3E) and MCF7-EXE (Fig. 3F) showed similar depletion of proliferative and MAPK signatures and enrichment of estrogen and tamoxifen response signatures. RAS signaling signatures were depleted with vandetanib treatment in all 3 cell lines (Fig. 3G). Similar transcriptional effects were seen in T47D and T47D-TAM (Fig. S4).

**Figure 3.**
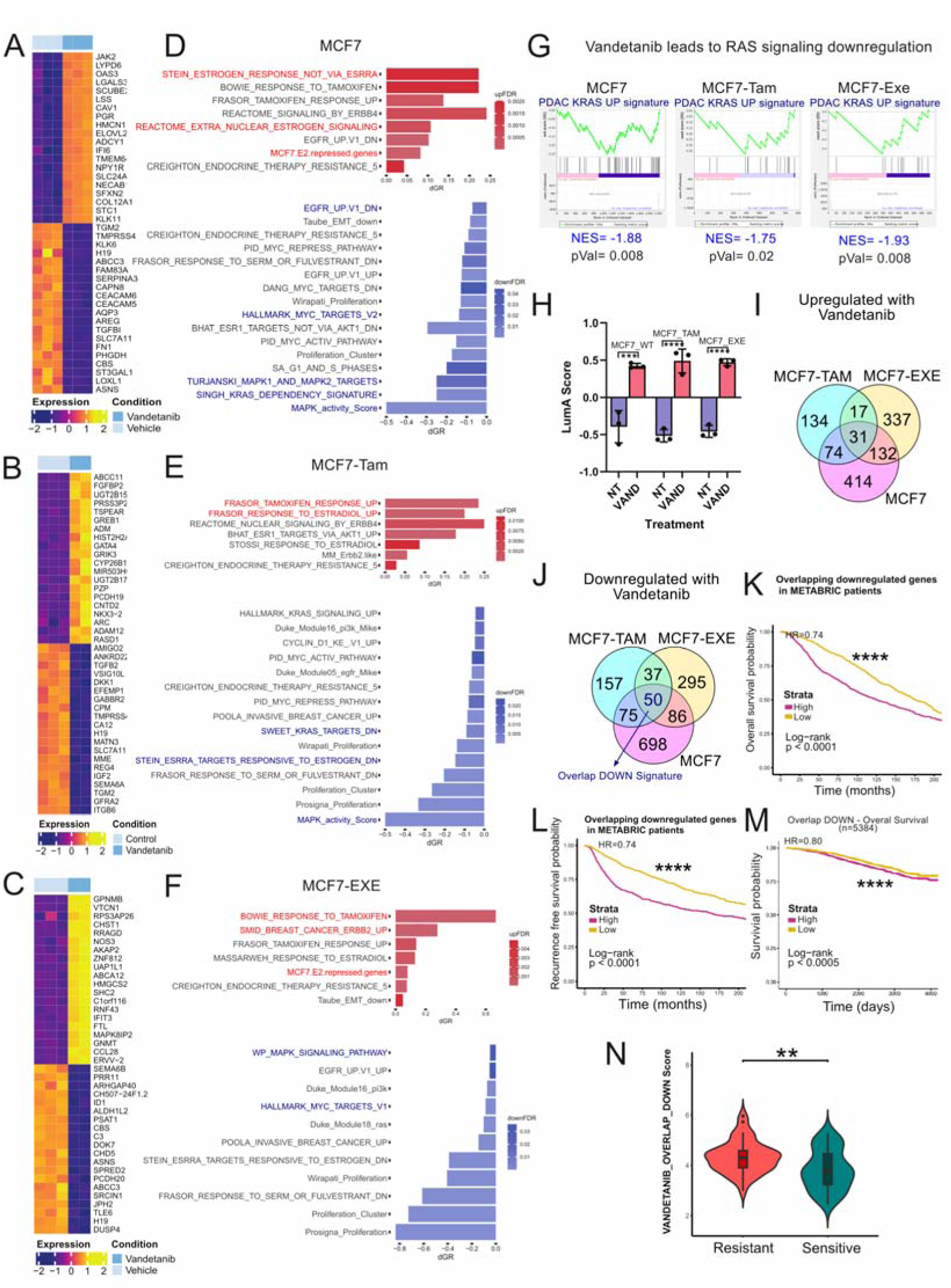
Vandetanib regulates gene expression resulting in a low proliferation, estrogen responsive Luminal-A like state. Heatmap and enrichment analysis of differentially expressed genes between vandetanib-treated vs control cells in (A) MCF7, (B) MCF7-TAM, (C) MCF7-EXE. Enriched and dep eted gene signatures in (D) MCF7, (E) MCF7-TAM, and (F) MCF7-EXE. (G) KRAS activity signatures were depleted in all 3 cell lines by vandetanib treatment supporting on target effects. (H) Vandetanib treatment increases association with the more indolent, endocrine therapy responsive Luminal A subtype, measured by correlation to Luminal A defining genes (centroid). Overlap of up- (I) and down- regulated (J) genes by cell line. (K) Signatures of down regulated gene signature are associated with overall survival and recurrence free survival (L) in ER+ breast cancer patients in METARIC and in (M) SCAN-B studies. (N) The downregulated gene set is enriched in clinically ET-resistant ER+ tumors. *p<0.05, **p<0.01, ***p<0.001, ****p<0.0001 by t-test with Welch correction for Wilcoxon test.

Enriched estrogen response signatures and reduced proliferation are hallmarks of the ET responsive Luminal A (LumA) molecular subtype that is associated with the most favorable patient outcomes(44). We analyzed the effect of vandetanib on genes that define the LumA subtype on the PAM50 assay (LumA centroid). This analysis demonstrated that vandetanib treatment resulted in a stronger LumA correlation supporting that treatment induced a more LumA-like indolent and ET responsive transcriptional phenotype (Fig. 3H).

To elucidate conserved molecular actions of vandetanib we determined overlap across the parental and ET-resistant MCF7 cell lines (Fig. 2I). Notably, significant commonly upregulated genes, including *SELENBP1* and *BAG1*, which are associated with reduced proliferation and improved survival in ER+ breast tumors (45-47). In the common downregulated genes, *AURKA* is associated with poor survival in ER+ breast cancer patients (48). Reduced expression of *ECT2* is associated with abrogation of MAPK signaling (49). Finally, several key regulators of cell cycle progression associated with worse outcomes and poor response to endocrine therapy in ER+ breast cancer are downregulated, including *CCND1*, *TOP2A*, and *CCNB1* (Fig. 2J).

To test if the conserved vandetanib down-regulated gene set is associated with prognosis in patients with ER+/HER2- breast cancer treated with endocrine therapy, we obtained survival annotated transcriptional data from METABRIC(50) and SCAN-B(51). Tumors were stratified by gene set score using median thresholding. METABRIC patients with low expression of genes that were downregulated by vandetanib had significantly improved OS (HR=0.75, p<0.0001, Fig. 3K) and RFS (HR=0.60, p<0.0001, Fig. 3L). Similarly, SCAN-B ER+ breast cancer patients with low expression of vandetanib downregulated genes demonstrated significantly lower mortality compared to high signature score (HR=0.87, p=0.034) (Fig. 3M). We next obtained gene expression data from patients with ER+/HER2- breast cancer profiled on ET and clinically annotated as sensitive or resistant (12). Resistant tumors had significantly higher expression of the vandetanib downregulated signature compared to sensitive tumors (4.3 vs 3.8, p<0.001, Fig. 3N). Cumulatively, these findings support that in ER+ breast cancer, vandetanib treatment promotes gene expression changes associated with lower proliferation, increased ET response, and improved survival.

### Vandetanib Treatment Enriches ER**α** Chromatin Occupancy at E2 Response Genes

Because vandetanib treatment resulted in enriched estrogen response signatures, we performed CUT&RUN to evaluate how vandetanib alters ERα chromatin binding. We performed peak differential analysis followed by gene peak annotation, comparing control and vandetanib treated cells. Taking the annotated peaks for each gene, we calculated the differential mean peak binding signals for gene signatures of interest. As expected, we found that vandetanib treatment increases binding of ERα to genes associated with estrogen activity, as well as genes associated with PI3K, STAT2 and ERBB2 signaling with vandetanib treatment (Fig. 4A). Genes with depleted ERα binding are associated with cell cycle transition, TNF and NOTCH pathway, among others.

**Figure 4.**
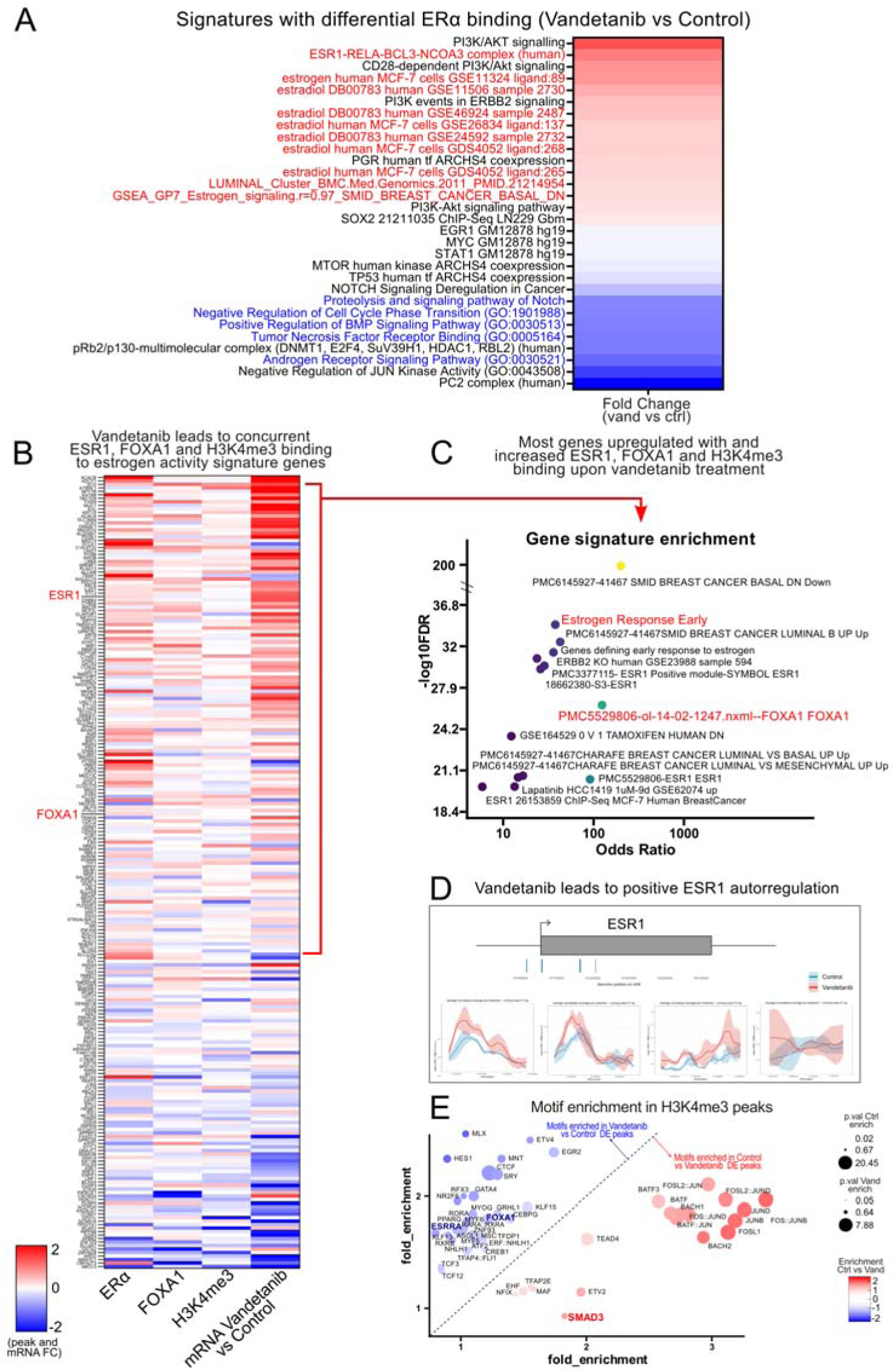
Enriched estrogen response signatures with vandetanib treatment correlate with enriched ERα binding site occupancy. (A) Heatmap showing gene expression signatures with enriched and depleted ERα promotor binding site occupancy. (B) Heatmap of estrogen response genes graphically demonstrating that changes in gene expression with vandetanib treatment correspond with changes in ERα promoter binding. Genes with increased expression also show enriched FOXA1 occupancy and promoter activity (H3K4me3). (C) GSEA for enriched estrogen activity genes demonstrate canonical E2 response signatures (D) Vandetanib results in increased ER binding to its own promotor representing a forward driven feedback mechanism. (E) H3K4me3 peaks filtered for known transcription driving promoter binding sites demonstrate enriched ER and FOXA1 binding motifs in upregulated sites demonstrating a central role for these transcription factors in changes in gene expression with vandetanib treatment.

To further characterize the changes on ERα occupancy induced by vandetanib, we also assessed chromatin occupancy of FOXA1, a pioneer factor that mediates ERα binding to chromatin sites and ERα co-localizing protein, and for H3K4me3, a histone mark that characterizes transcriptional active sites and active promotor regulatory elements. FOXA1 showed increased occupancy to genes containing ERα binding sites but decreased occupancy to gene signatures indicating ERα activity in vandetanib-treated cells (Fig. S5A) and this was similar for H3K4me3 marks (Fig. S5B).

To assess the interaction of ERα, FOXA1 and H3K4me3 and the effect on expression of ERα-response genes, we analyzed the concurrent peak occupancy with mRNA expression in vandetanib-treated cells versus control (Fig. 4B). Generally, genes that gain ERα occupancy also gain FOXA1 and H3K4me3 occupancy, and this is associated with increased gene expression. GSEA of the genes with increased ERα, FOXA1 and H3K4me3 binding and transcriptional activity on vandetanib-treated cells showed these genes are associated with estrogen response (Fig. 4C). This reinforces that vandetanib leads to enriched ERα and FOXA1 chromatin activity on key estrogen response genes. Furthermore, we found increased ERα binding to ESR1 transcriptional start site and gene body peaks, indicating that vandetanib promotes a positive auto-regulatory loop in ESR1 expression (Fig. 4D). We next looked at enriched H3K4me3 sites with vandetanib treatment filtering for known promotor activating loci to assess transcription factor motifs enriched at those sites. Both ER and FOXA1 motifs were enriched at enhancer sites supporting a central role in those proteins mediating the gene regulatory effects of vandetanib (Fig. 4E).

### Functional Signatures of Vandetanib Response

Using multiplex inhibitor bead enrichment coupled to mass spectrometry (MIB/MS), we investigated the reprogramming of kinase signaling networks in response to vandetanib (38). In this technique, kinases are enriched by incubating cell extracts with type 1 kinase inhibitors immobilized on Sepharose beads (Fig. 5A). Vandetanib treatment caused depletion of known vandetanib targets (52) in all 3 models (Fig. 5B), despite large baseline differences in activity, (Fig. 5C). Similar results were seen for T47D and T47D-TAM (Fig. S6).

**Figure 5.**
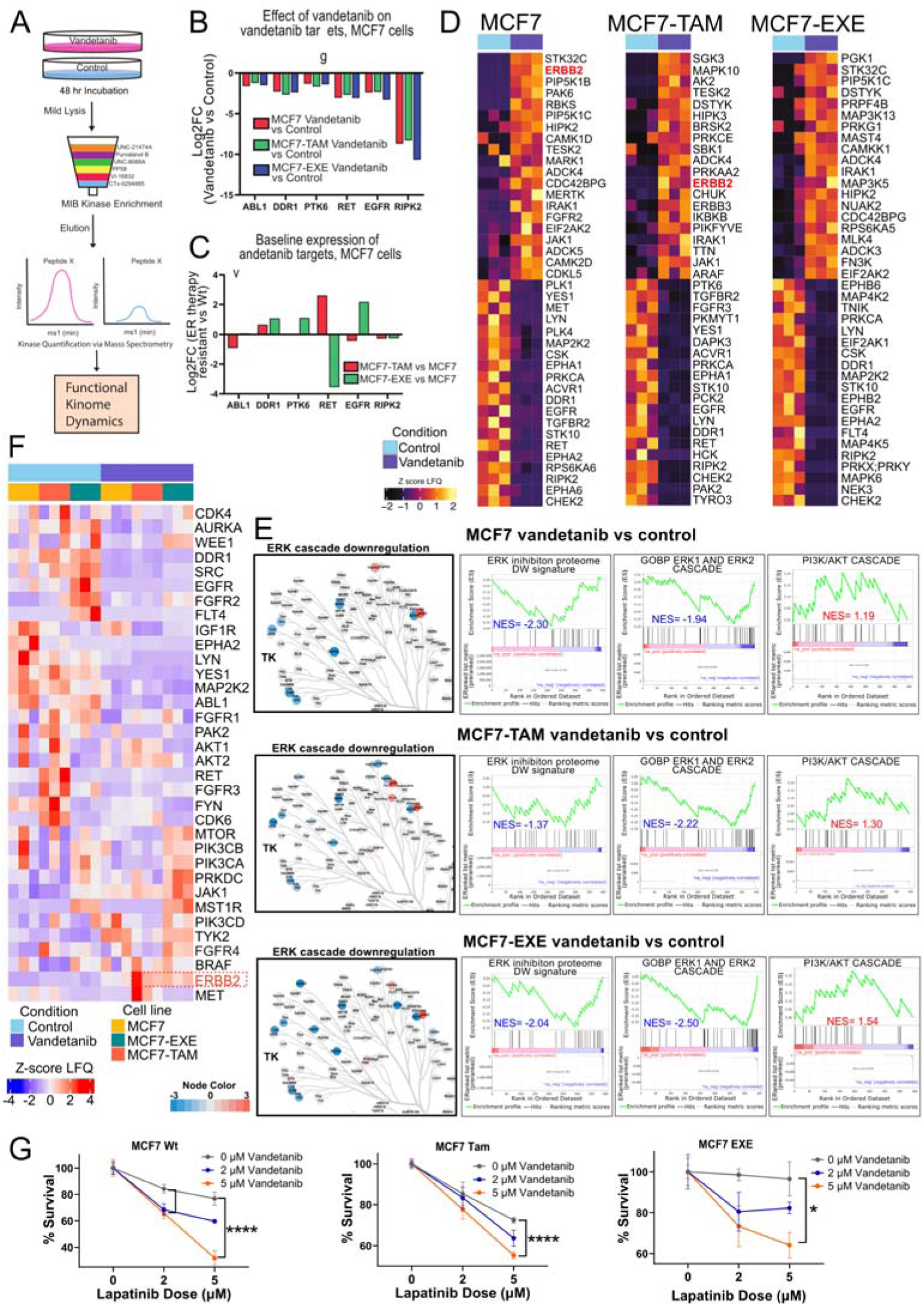
Functional Changes in Protein Signaling Networks with Vandetanib Treatment Identifies Targetable Adaptive Resistance Via Upregulation of HER2 activity. (A) Multiplex inhibitor bead mass spectrometry assay workflow. (B) Vandetanib resulted in depletion of known vandetanib targets with similar magnitude observed across cell lines. (C) Baseline expression of vandetanib targets in tamoxifen and exemestane resistant cell lines compared to WT MCF7. (D) Heatmap showing enriched and dep eted proteins after vandetanib treatment. (E) ERK pathway proteins were downregulated and PI3K upregulated with vandetanib treatment in all 3 cell lines. (F) Heatmap showing enrichment levels of druggable kinases of sensitive and resistant MCF7 cell lines before and after vandetanib treatment. Of note, ERBB2 (encoding HER2) was enriched in all 3 cell lines after treatment and RET was depleted in MCF7 and MCF7-TAM. (G) Sensitivity plots of MCF7 cells co-treated with vandetanib and HER2 inhibitor lapatinib demonstrating enhanced sensitivity with vandetanib treatment. *p<0.05, ****p<0.0001, t-test with Welch correction.

Enriched and depleted proteins by cell line are shown in Fig. 5D. Overlap analysis showed that kinome dynamics were heterogenous. Common upregulated proteins included AURKA, PLK1, PIP4K2A, which promote proliferation (53-55). Common depleted proteins with vandetanib treatment included proliferation, promoting cell cycle regulators CDK1, CDK2, and PKMYT1. Depletion of MAP2K2 (MEK2) supports downregulation of the MAPK signaling pathway in response to vandetanib treatment, consistent with Fig. 1A. Gene set enrichment analysis similarly demonstrated downregulation of the ERK cascade (56) with largely consistent depletion of specific node proteins (Fig. 5E). PI3K/AKT pathway and ERBB2 signaling were also upregulated. Interestingly these were two pathways identified with enriched ERα chromatin occupancy (Fig. 4A), demonstrating the transcriptional basis for protein signaling network adaptation.

### Inhibition of Adaptive Resistance Pathways Enhances Vandetanib Efficacy

Upregulated functional kinases may identify mechanisms of adaptive bypass resistance, so we assessed activity of druggable kinases (23) by treatment (Fig. 5F). We performed co-treatment assays on the principle that inhibiting mediators of adaptive resistance should enhance sensitivity to vandetanib. Because ERBB2/HER2 was enriched in all 3 MCF7 cell lines, we co-treated cells with vandetanib and the FDA approved HER2 small molecule inhibitor lapatinib. Treatment with lapatinib resulted in significantly augmented vandetanib response measured by reduction in cell viability in all 3 cell lines, (Fig. 5G), validating increased reliance on HER2-activity in vandetanib treated cells.

### Vandetanib Response at Single Cell Resolution in Primary ER+ Breast Cancer

We used our previously validated primary tumor *ex vivo* short-term drug treatment platform coupled to single-cell RNA sequencing (40), to assess how primary ER+ breast tumor cells respond to vandetanib treatment. Tumor cells were treated with control or vandetanib containing media for 12 hours and assayed by scRNAseq. Uniform manifold projection (UMAP) was performed using Seurat v3 (57). Cell populations were annotated using known transcriptional profiles and canonical markers (Fig. 6A) and by treatment group for tumor cells (Fig. 6B). Malignant (tumor) cells were distinguished from non-tumor epithelial cells as cells having significant inferred copy number alterations by InferCNV (58). Differentially expressed genes were identified using a false discovery rate (FDR) cutoff of < 0.05 in the iCNV+ tumor cell compartment with vandetanib treatment compared to control (Fig. 6C). Enriched and depleted pathways in vandetanib treated tumor cells relative to control treated were determined (Fig. 6D). As was seen in cell lines, estrogen response genes and signatures associated with tamoxifen sensitivity were enriched.

**Figure 6.**
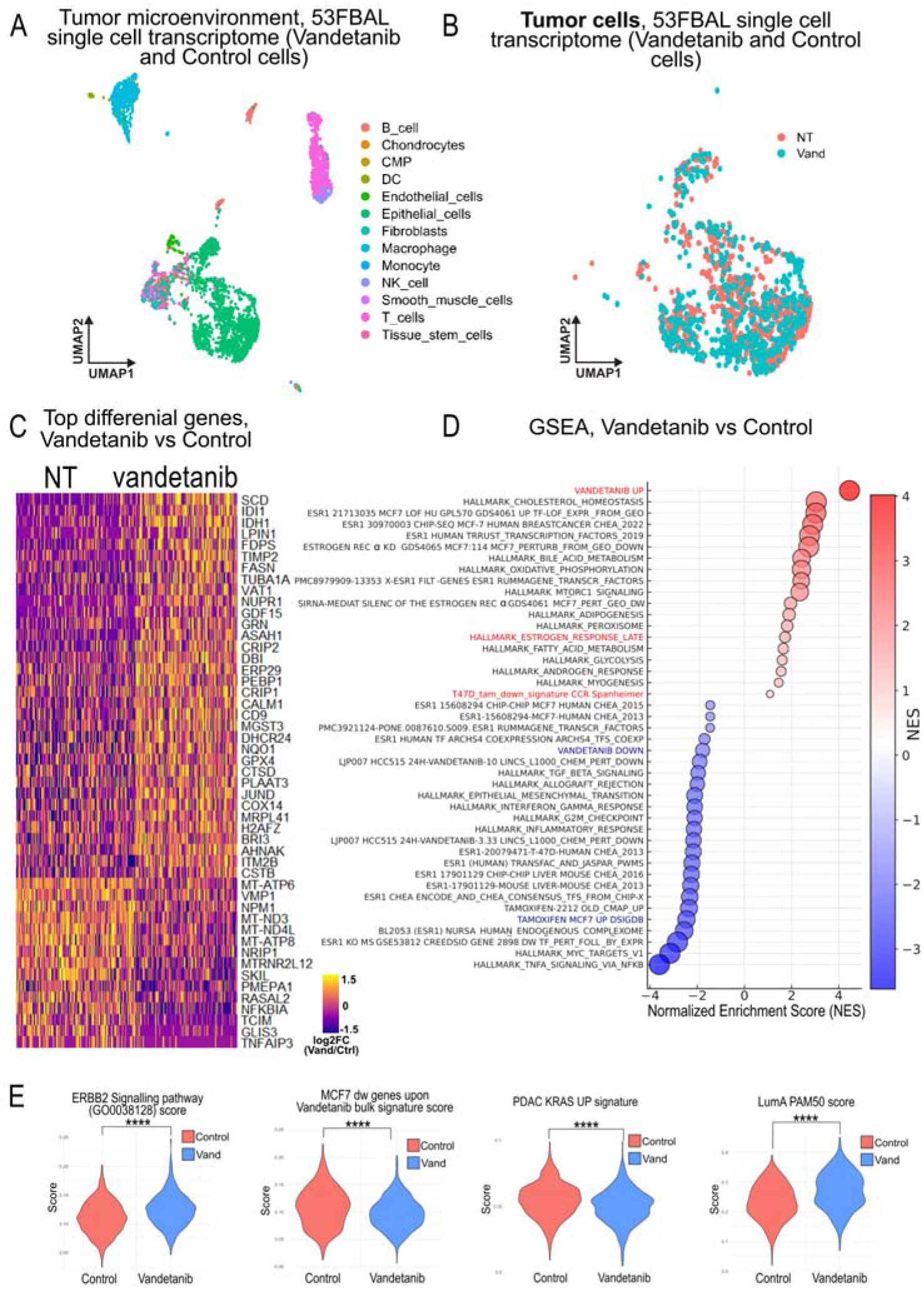
Primary ER+/HER2- breast tumor cells demonstrate conserved transcriptional effects with vandetanib treatment. (A) Single-cell RNA sequencing uniform manifold projection (UMAP) annotated by cell type within a primary ER+ human breast tumor. (B) Tumor cells were computationally isolated as having significantly inferred copy number changes and plotted by treatment condition. (C) Heatmap showing differentially expressed genes between vandetanib treated vs non-treated tumor cells. (D) Enriched and depleted pathways of vandetanib treated vs non-treated tumor cells. (E) ERBB2 activity signature score was increased in vandetanib treated tumor cells as seen in cell lines. Down regulated genes in vandetanib treated cell lines are depleted in vandetanib treated primary tumor cells as were KRAS activity signature demonstrating on target effects. LumA correlation increased in vandetanib treated tumor cells as was seen in cell lines. Significant genes filtered by adjusted p value <0.05 (Wilcoxon test) and significant signatures filtered by < 0.1 FDR on GSEA analysis.

To demonstrate the in-human relevance of the cell line findings, we examined salient features seen in cell lines in these primary tumor cells. First, we tested the overlapping downregulated genes after vandetanib treatment across the 3 MCF7 cell lines (Fig. 3J) and these genes were lower in vandetanib treated primary tumor cells (Fig. 6E). Depletion of KRAS signature supports on-target effects. Next, LumA correlation score was assigned to each tumor cell using a validated single cell PAM50 classifier (59) and treated tumor cells had a stronger correlation to LumA. Thus, as with cell lines, short-term vandetanib treatment alters the transcriptional profile of ER+/HER2- primary tumor cells towards a more indolent and estrogen responsive phenotype. Lastly, ERBB2 signaling was enriched supporting that primary human tumor cells also induce vulnerability to HER2 inhibitors in response to vandetanib treatment.

To explore features of vandetanib response we used non-negative matrix factorization (NMF) (Fig. S7A). This analysis yielded metagene modules that were enriched in vandetanib treated samples (module 1) and depleted in vandetanib treated cells (Module 6) (Fig. S7B). We next identified features that may be associated with vandetanib response by comparing modules 1 and 6 gene expression profiles. Module 1 (vandetanib resistant) had expected high expression of genes that increase with vandetanib treatment, consistent with poor response. Module 6 was enriched for MYC targets, which is known to support tamoxifen resistance, but may support a transcriptional profile of vandetanib sensitivity (Fig. S7C). Interestingly, module 1 was enriched for hallmark estrogen response genes and our previously generated single cell tamoxifen response score (40). Therefore, cells with poor vandetanib response may have enhanced tamoxifen sensitivity (Fig. S7D) supporting dual treatment to target distinct cell types within the tumor. Lastly, we looked at our human tumor derived signatures of high GDNF/RET expressing human ER+-breast tumors (22). Genes downregulated in GDNF/RET high human tumors had low expression in vandetanib sensitive cells, supporting RET activity as a key feature of responsive cells (Fig. S7E).

### Vandetanib in ER+/HER2- Breast Cancer Patient Derived Xenografts

Mice implanted with ER+/HER2- patient derived xenografts were treated with control or vandetanib gavage. Tumor FFPE sections showed expected morphology and expression of ER and lack of HER2 expression (Fig. 7A). For HCI011 vandetanib significantly reduced tumor growth at 6 weeks (Fig. 7B). HCI05 tumors had no reduction in growth with vandetanib, demonstrating a vandetanib sensitive (HCI011) and resistant (HCI05) model. As expected, the sensitive model (HCI011) had higher expression of genes associated with RET activity in human ER+ tumors, supporting that RET activity is also a key feature of vandetanib response in PDX (Fig. 7C). Further, high expression of the vandetanib downregulated genes in MCF7 was also seen in HCI011. Lastly, NMF1 module (Fig. S7), which are genes highly expressed in vandetanib resistant human tumor cells were expressed more highly in the resistant HCI05 PDX.

**Figure 7.**
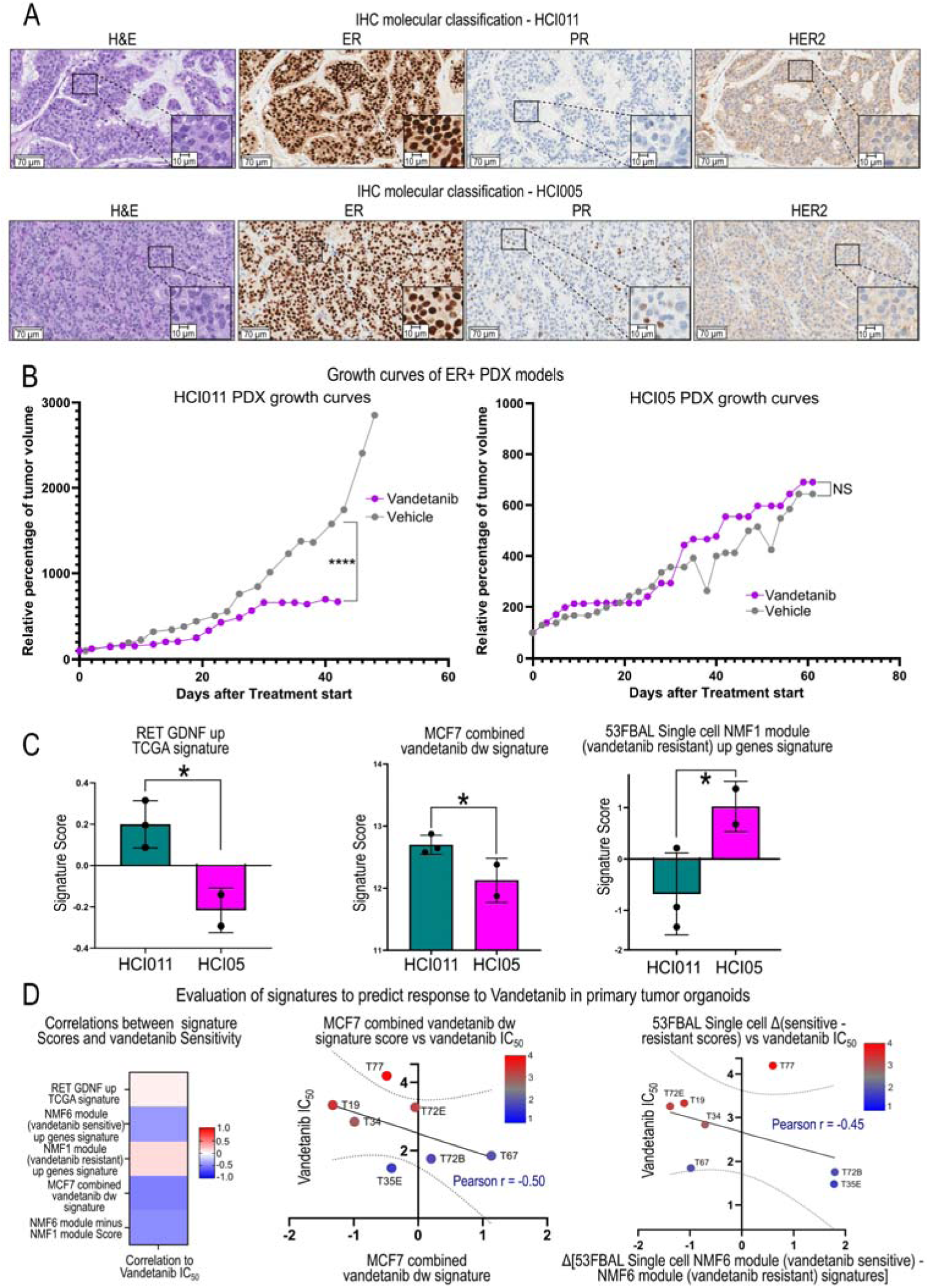
Vandetanib signatures predict sensitivity in patient derived models. (A) H+E slides demonstrate histology (H+E) and strongly ER+, weakly PR+, and HER2- status for two PDX models. (B) Five mice for each PDX were randomized to treatment with control or vandetanib (50 mg/kg daily) by oral gavage. HCI011 had reduced growth with vandetanib and HCI05 did not. ****p<0.0001 by Linear mixed-effects model growth slope test after 20 days post treatment (C) The vandetanib sensitive HCI011 had higher baseline expression of vandetanib down-regulated genes and the vandetanib resistant HCI05 had higher expression of our primary tumor NMF6 derived resistance signature, supporting the utility of these signatures to identify tumors that are responsive or resistant to vandetanib. *p<0.05 t-test with Welch correction. (D) PDO signature correlation with response to vandetanib measured by IC50 and vandetanib-tamoxifen synergy. A score of the difference between our human tumor NMF derived vandetanib resistance and vandetanib sensitive signatures most strongly correlated with response to vandetanib (high score indicated resistance (high IC50)).

### Vandetanib Response Signatures in PDX and PDO

We next used our PDO mRNA sequencing and sensitivity data (IC50) to test performance of derived signatures on predicting vandetanib sensitivity in human tumor derived models. We tested if 1) our TCGA human tumor derived signature of RET activity (GDNF/RET up TCGA Signature) (22), 2) cell line derived signature of down-regulated genes with vandetanib treatment in MCF7 (Fig. 3J)., and 3) a single cell human tumor vandetanib response score of the difference between sensitive cell genes (NMF6) and resistant cell genes (NMF1) (Fig. S7B/C) were associated with sensitivity to vandetanib. We found that both the MCF7 down signature and the human tumor single cell response score were inversely correlated with IC50 of ER+ PDO demonstrating that a high signature score is associated with more sensitive PDO (Fig. 7D).

## DISCUSSION

Resistance to endocrine therapy (ET) is a significant driver of poor patient outcomes, despite advances in understanding how ER+ breast tumors respond to therapy. The identification of MAPK pathway activation as a driver of resistance to ET has generated significant interest in targeting this pathway to enhance ET response, but this has not translated to the clinic. This highlights the need for clinically actionable kinase targeting strategies in MAPK-activated, ET-resistant breast tumors and biomarkers of patients most likely to respond.

Kinase signaling networks can be plastic and inhibiting one node can lead to adaptive bypass through another pathway. We identified PI3K and HER2 as potential mediators of adaptive resistance to vandetanib treatment, and targeting HER2 with lapatinib enhanced sensitivity to vandetanib. ERα occupancy was enriched at genes mediating PI3K and ERBB2 signaling upon vandetanib treatment, suggesting that ER rewiring may contribute to the adaptive kinase landscape. Activating HER2 could open a number of efficacious therapeutic options for patients. While HER2 activity was increased, there was no change in HER2 RNA expression, highlighting the need for protein activity level assessment to identify rationale cotreatments. To this end, our group has recently developed a method for comprehensive-kinome characterization on limited sample, such as a core needle biopsy, which could enable on-treatment assessment of adaptive responses and facilitate real-time, patient specific interactive cotreatment regimen design (60).

RET mediates effects of vandetanib and we found that RET-regulated genes are a key marker of response to vandetanib in patient derived models, but CRISPR knockout of RET unexpectedly resulted in reduced, rather than enhanced, sensitivity to tamoxifen. ER expression was reduced after RET knockout, as were ER induced genes, and this likely accounts for decreased sensitivity to tamoxifen. While ER has 2 binding sites in the RET promoter that activate RET expression (61), no reciprocal mechanism by which RET regulates ER is known. RET expression and ER-activity are correlated, RET knockout may select for cells that have lower baseline ER-activity by enriching populations that are less dependent on RET signaling. Further studies are needed to model emerging resistance to RET and ER inhibitors and to track relative dependencies on RET and ER within emerging resistant tumor cell subpopulations.

Our human tumor single cell analysis reinforces key *in vitro* findings. RET activity is a key component sensitivity to vandetanib in ER+ breast tumors. Cells categorized within the vandetanib sensitive metagene module have low expression of genes that are downregulated in GDNF/RET high human tumors, indicating that vandetanib sensitivity is associated with our previously described high RET-activity phenotype (22). Importantly, we show that vandetanib response is not homogeneous within tumors. Instead, distinct sensitive and resistant populations can be resolved at single-cell resolution. Resistant cells express features associated with tamoxifen response, consistent with a conceptual model where the population that persists under vandetanib remains sensitive to ET. In contrast, a vandetanib sensitive cells were enriched for MYC targets and programs linked to tamoxifen resistance, consistent with a vandetanib-sensitive but ET-refractory state. These results indicate that within heterogenous human tumors, vandetanib preferentially eliminates a MYC-high, ET-poorly responsive compartment, while a more estrogen-responsive, tamoxifen-sensitive compartment persists.

Similarly in PDX we identified a vandetanib-sensitive and a resistant ER+/HER2− tumor. The sensitive PDX expressed a RET activity signature consistent with high RET-activity phenotype, as well as the vandetanib-sensitivity gene programs identified in cell lines, whereas genes defining the vandetanib-resistant human tumor cells were more highly expressed in the resistant model. These results show that the same vandetanib-linked transcriptional states identified in cell lines and primary tumor cells stratify response in orthotopic PDX models, indicating that these states reflect true biological vulnerabilities rather than model-specific features, and that these signatures could be used to as to predict response to vandetanib.

In summary, we demonstrate that ER+ breast cancer cells become more sensitive to ET with vandetanib treatment. Vandetanib enriches ER binding site occupancy at canonical E2 response genes altering gene expression and kinase signaling resulting in a lower proliferative, more estrogen-responsive luminal A-like state with an induced vulnerability to HER2 inhibitors. We define response signatures that strongly correlate with sensitive and resistant patient derived organoid and xenograft models. These results support that vandetanib treatment could benefit select patients with ER+ breast cancer and that future trials should evaluate response signatures as predictive biomarkers.

## ACKNOWLEDGEMENTS

We would like to thank Dr Alana Welm at the University of Utah for generously providing the patient derived xenograft models.

## Author Contributions

Conceptualization: RTK, GLJ, PMS

Methodology: AAW, ECB, DOO, MPE, ASB, JRR

Investigation: AAW, RTK, SH, CHNCT, ATS, DOO, MPE, ASB, JRR

Visualization: SH, ECB, JRR

Funding Acquisition: LAC, CMP, PMS

Supervision: GLJ, PMS

Writing – original draft: RTK, SH, PMS

Writing – review and editing: all authors

## Data and Materials Availability

Single cell and bulk RNA sequencing data was made available at the Sequence Read Archive (SRA) under BioProject PRJNA1177387. Mass spectrometry protein sequencing results are included in this manuscript as Supplementary Table 1. All original code has been deposited and is publicly available in the Github repository https://github.com/santiagohaase/Spanheimer-lab-Kinase-Plasticity-in-Response-to-Vandetanib-Enhances-Sensitivity-to-Tamoxifen

