## Supplemental figures for "Kinase Plasticity with Vandetanib Treatment Enhances Sensitivity to Tamoxifen in Estrogen Receptor Positive Breast Cancer"

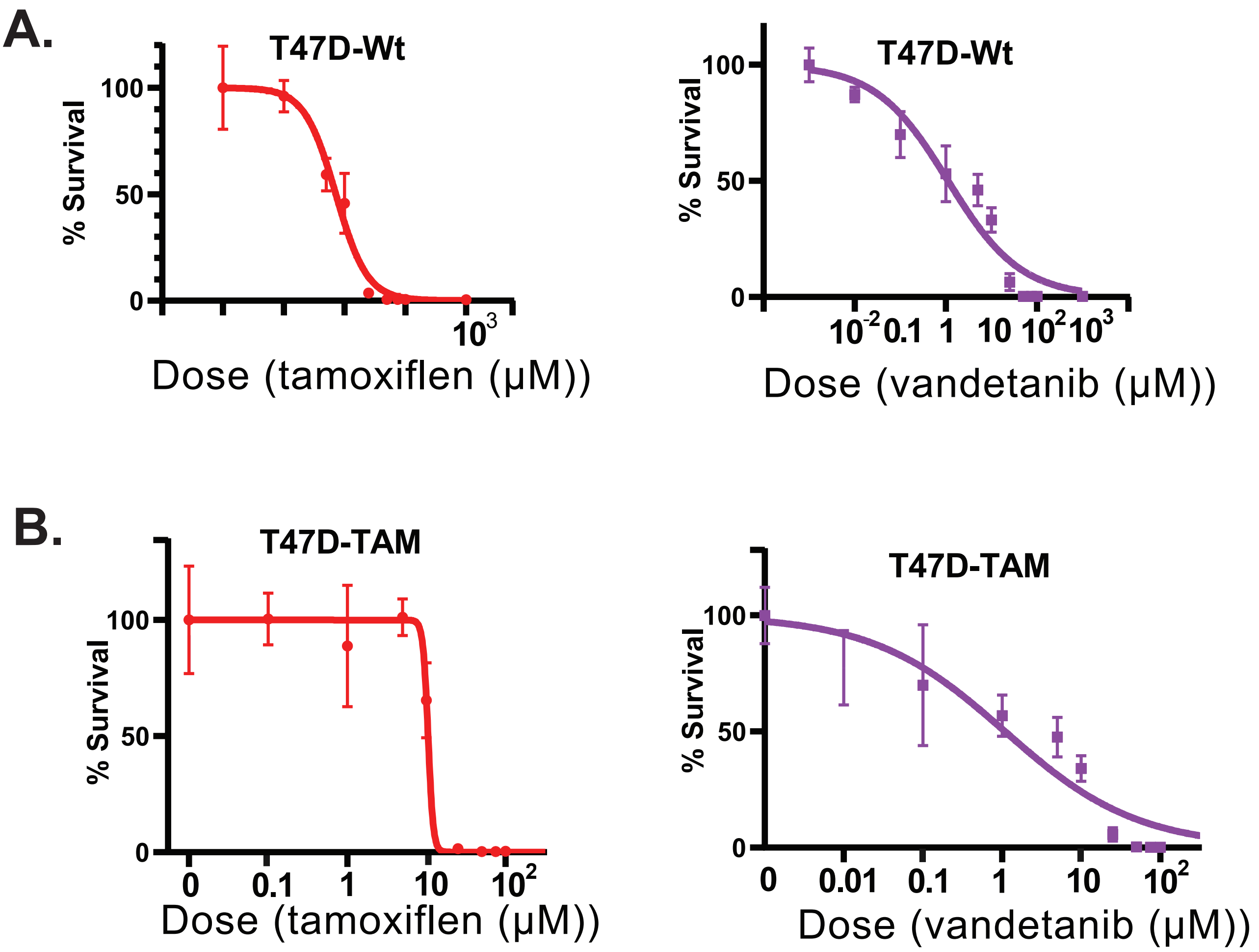


Supplemental Figure 1: Tamoxifen and vandetanib sensitivity curves in T47D (A) and T47D-TAM (B).

A

| Label | Pt Age At Surgery | Gender | Race | Primary Diagnosis | Overall grade at Path | ER% | PR% | Her2 Ihc | pT | pN | PAM50 |
|---|---|---|---|---|---|---|---|---|---|---|---|
| T19 | 58 | Female | Black | IDC | 3 | 99 | 9 | 2 | 2 | 0 | LUMA |
| T34 | 50 | Female | Black | IDC | 1 | 90 | 70 | 1 | 1 | 0 | LUMA |
| T35 | 49 | Female | Other | Mixed Ductal | 1 | 95 | 100 | 1 | 1 | 1 | LUMB |
| T67 | 57 | Female | White | IDC | 2 | 100 | 95 | 0 | 2 | 1mic | HER2 |
| T72 | 36 | Male | White | IDC | 3 | 95 | 70 | 1 | 2 | 1mic | Basal |
| T77 | 74 | Male | White | IDC | 3 | 95 | 20 | 1 | 1 | 0 | Basal |
| T4 | 47 | Female | Black | IDC | 3 | 0 | 0 | 0 | 3 | 1 | Basal |

IDC= Invasive Ductal Carcinoma - pT =Pathologic Tumor Stage - pN = Pathologic Nodal Stage

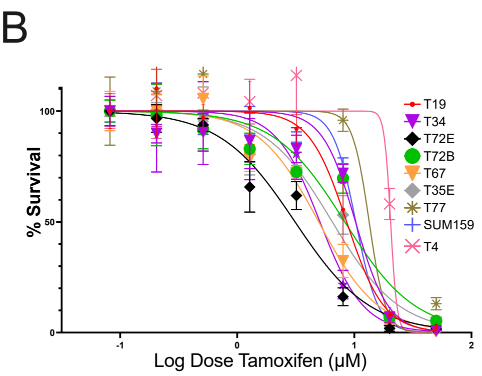


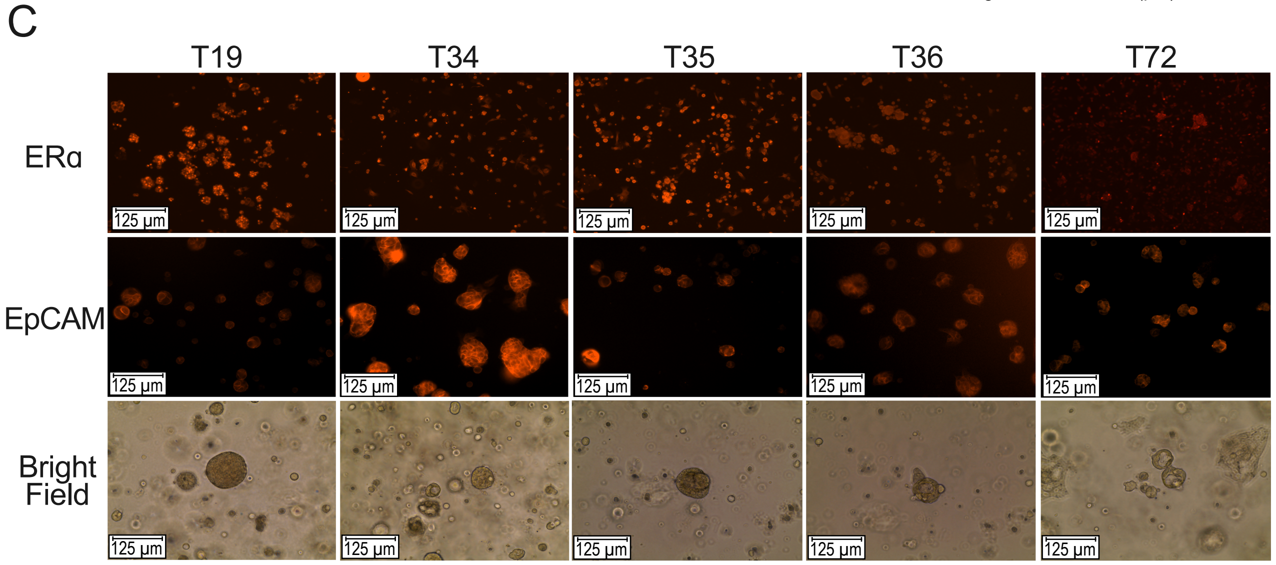


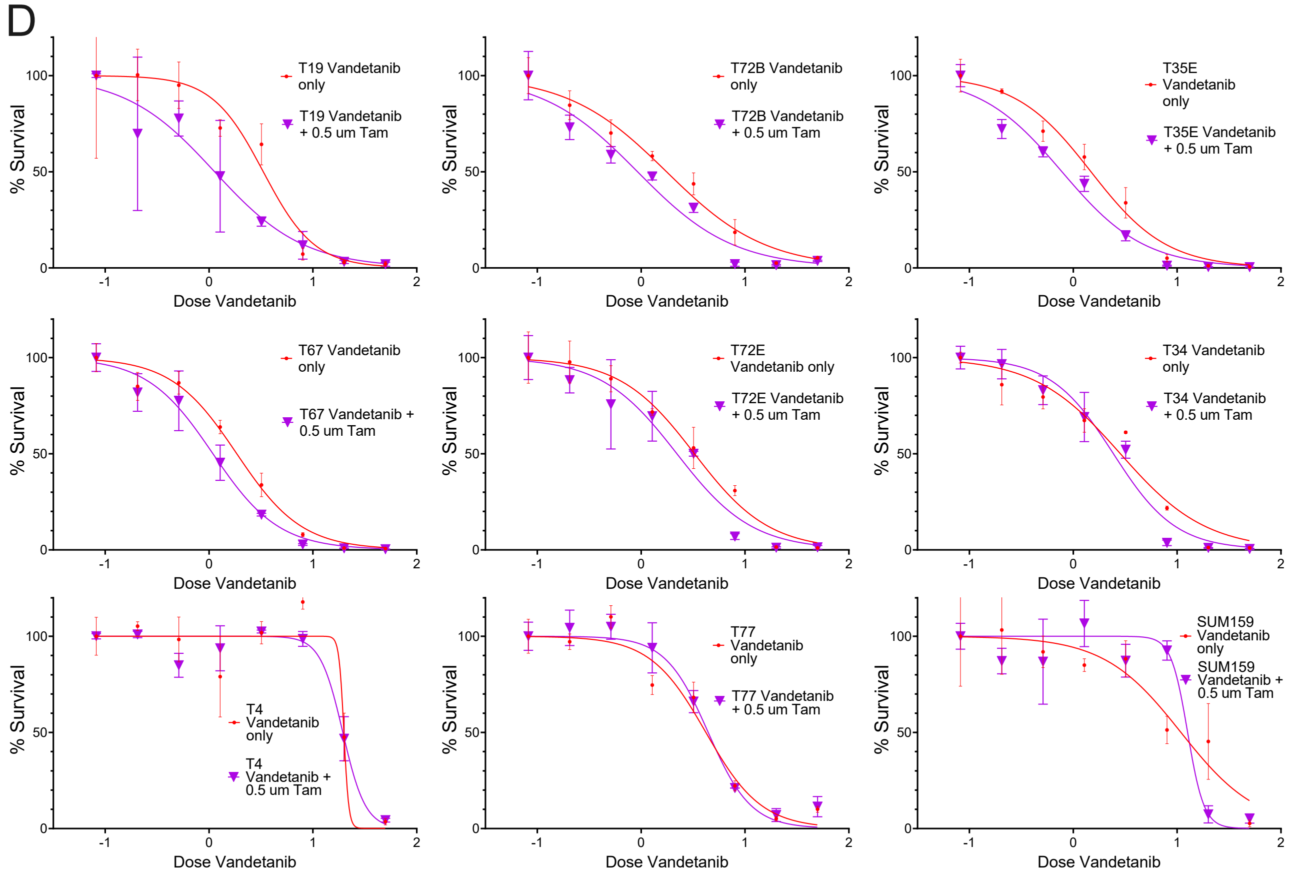


Supplemental Figure 2: Clinical data and molecular classification of patient-derived breast cancer organoids (PDOs). B) PDOs dose-response curves to tamoxifen. C) PDOs immunofluorescence stainings for ERα and EpCAM (Epithelial Cell Adhesion Molecule). D) Side-to side dose-response curves for PDOs to vandetanib alone and to vandetanib combined with a fixed dose (0.5 µM) of tamoxifen. PDOs curves are arranged in decreasing order of synergy.

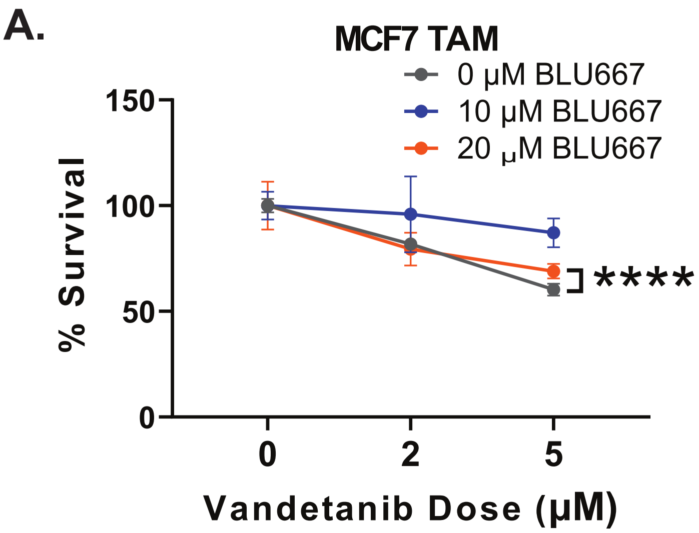


Supplemental Figure 3: Acute inhibition of RET with the selective inhibitor BLU667 reduced sensitivity to vandetanib in MCF7-TAM (A) but not in the low RET expressing MCF7-EXE (B). (****$p<0.0001$ by t-test with Welch correction.)

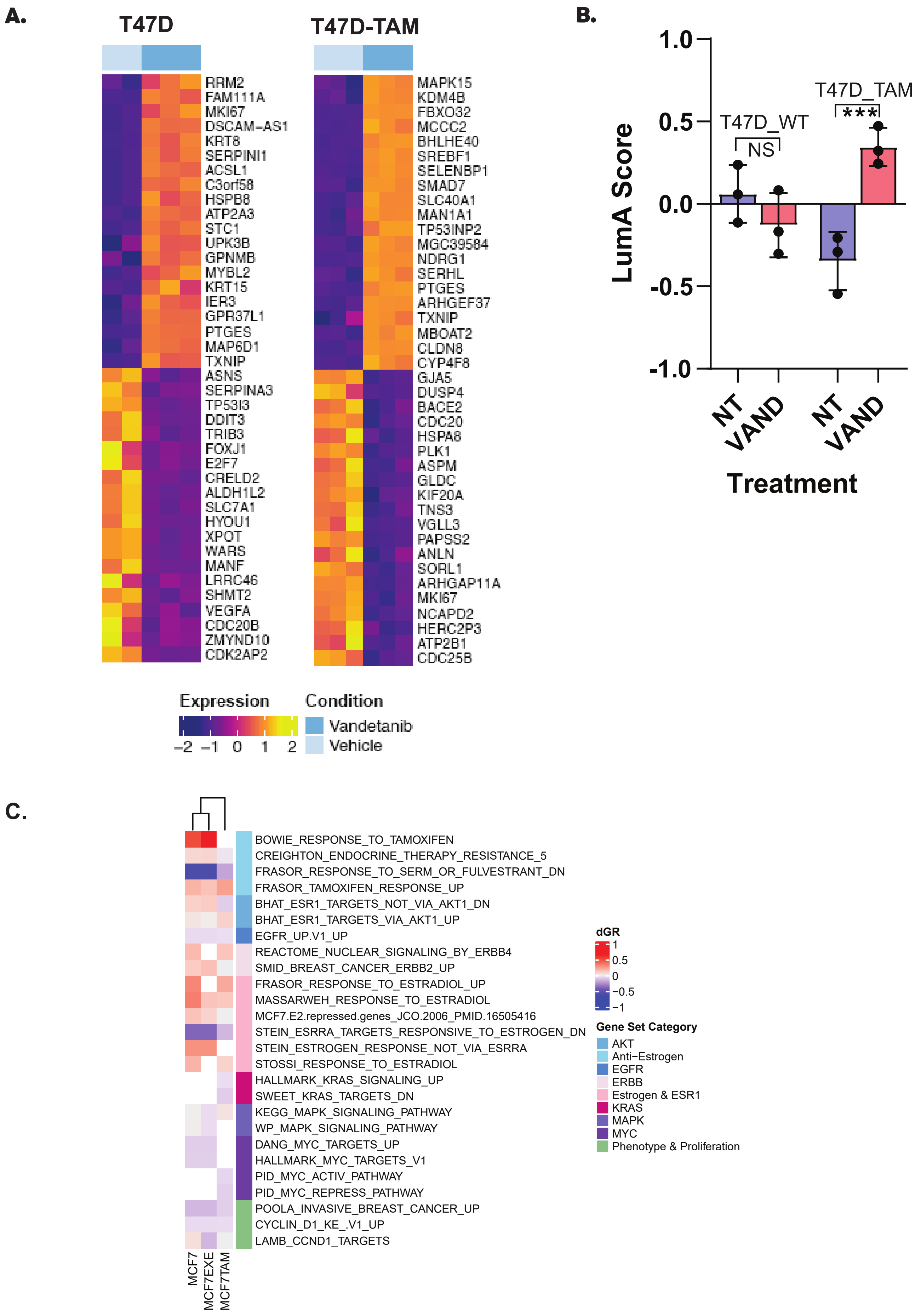


Supplemental Figure 4: (A) Heatmap showing up- and down-regulated genes in T47D and T47D-TAM (Vandetanib vs Control. (B) Vandetanib treatment increased luminal A correlation in T47D-TAM but not T47D. (C) Gene set enrichment shows enrichment of ER signaling pathways after vandetanib treatment. (***p<0.005, Wilcoxon test.)

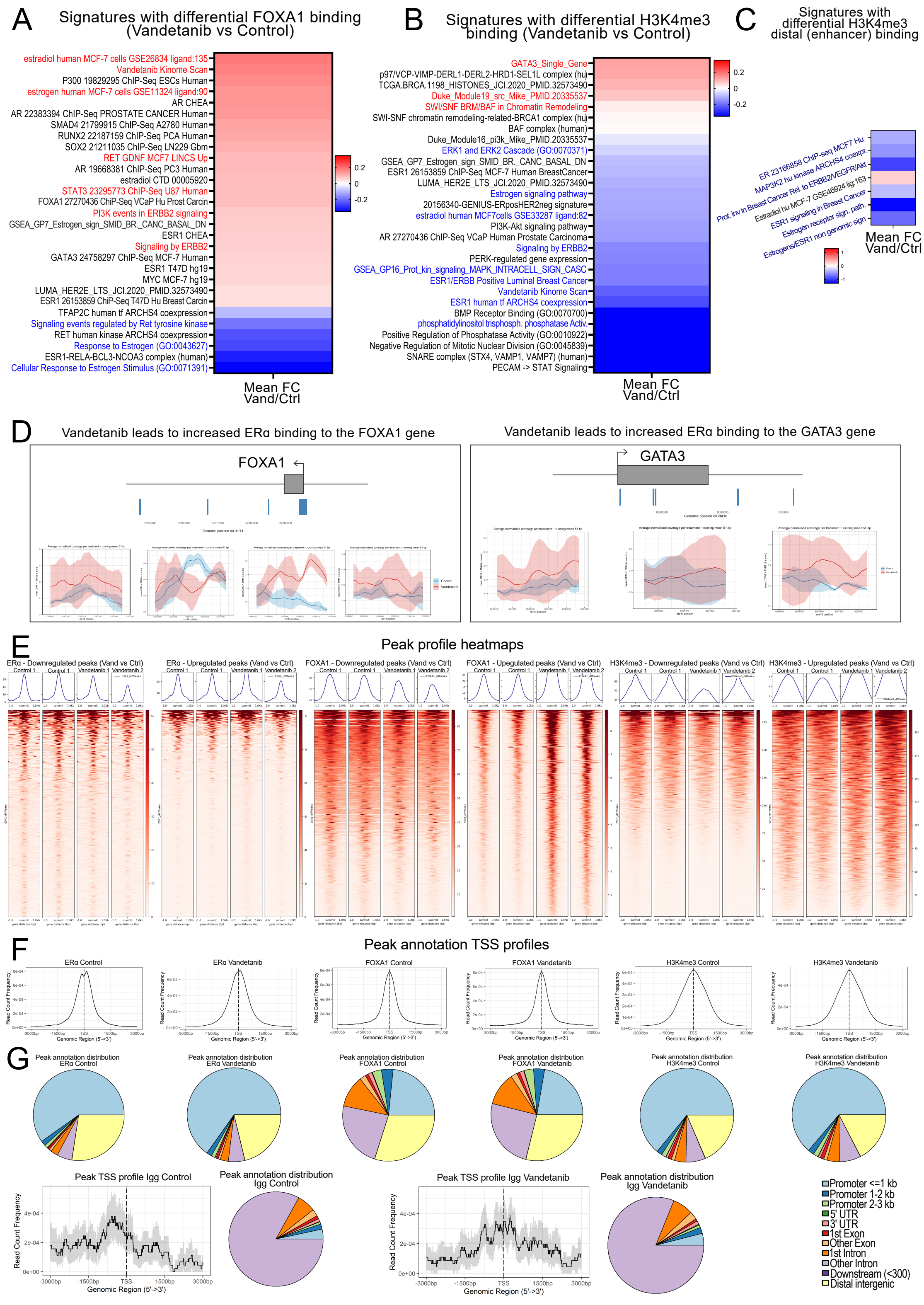


Supplemental Figure 5: (A) Heatmap showing selected gene expression signatures with enriched and depleted FOXA1 binding site occupancy. (B) Heatmap showing selected gene expression signatures with enriched and depleted H3K4me3 binding site occupancy. (C) Heatmap showing selected gene expression signatures with enriched and depleted H3K4me3, using only H3K4me3 peaks distal to genes. (D) Coverage plots of ERα occupancy to FOXA1 and GATA3 genes. (E) Peak profile heatmaps for ERα, FOXA1 and H3K4me3 differentially upregulated and downregulated (Vandetanib versus control) peaks. (F) Peak transcriptional start site (TSS) profiles for ERα, FOXA1 and H3K4me3 (Control and Vandetanib-treated) annotated peaks. (G) Peak gene location distribution for ERα, FOXA1 and H3K4me3 (Control and Vandetanib-treated) annotated peaks; and TSS and gene location profiles for Igg Cut and Run background controls.

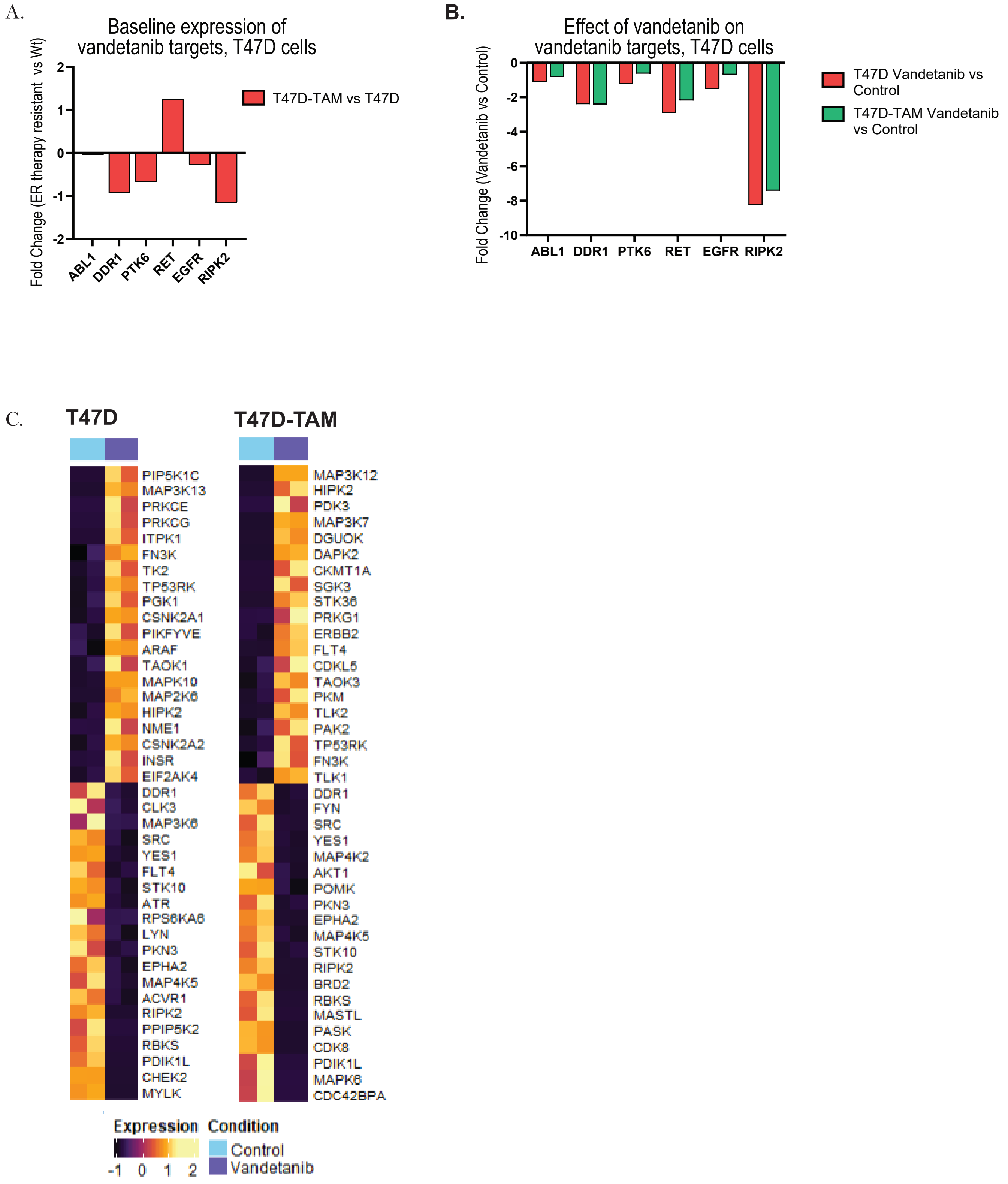


Supplemental Figure 6: (A) Baseline changes in vandetanib target genes in T47D-TAM vs T47D measured on MIB/MS. (B) Depletion of direct vandetanib targets in T47D and T47D-TAM. (C) Heatmap showing enriched and depleted proteins after vandetanib treatment.

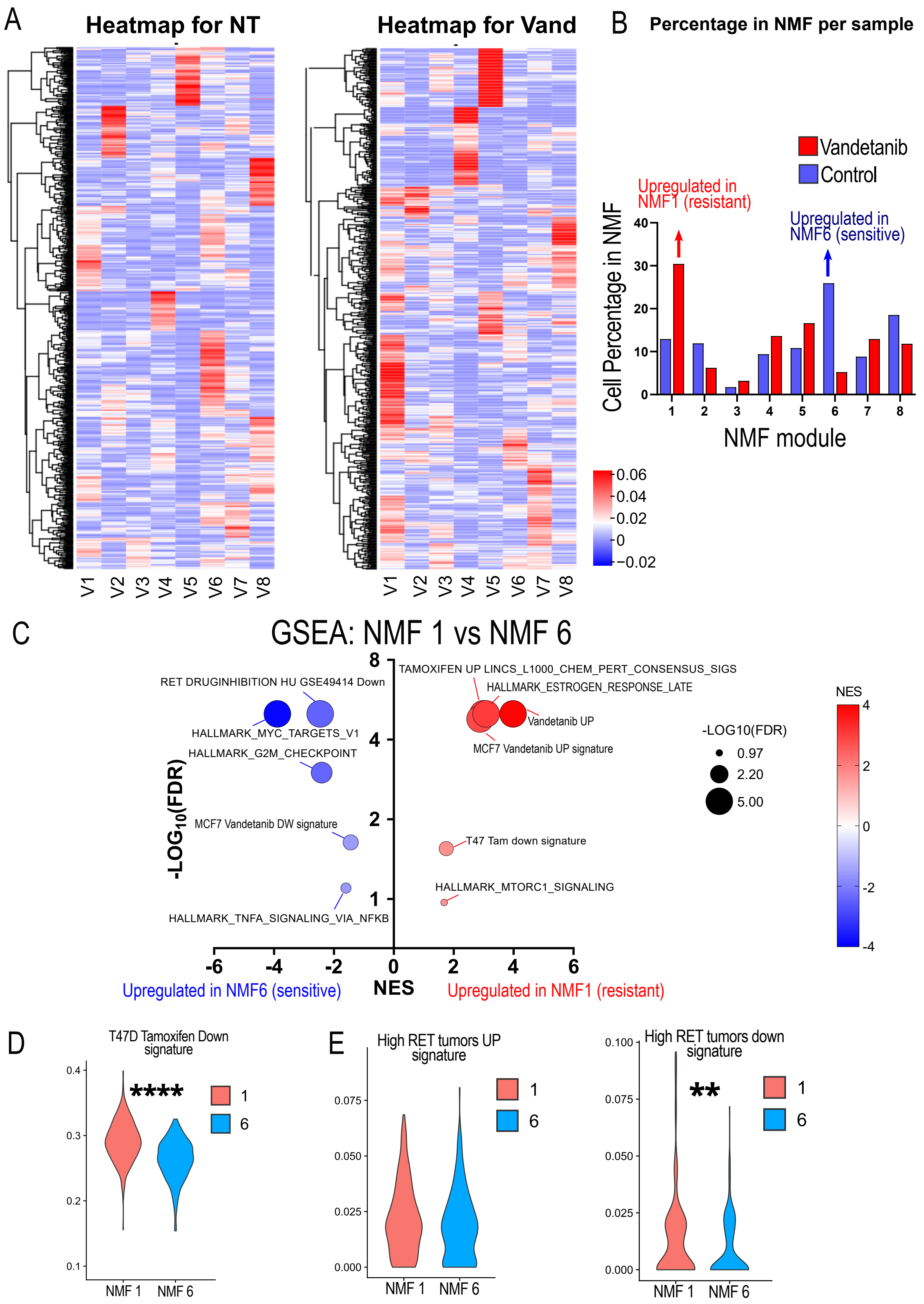


Supplemental Figure 7: (A) Non-negative Matrix factorization (NMF) Module Heatmap in ER negative breast cancer organoid tumor cells. (B) Cell percentage from treatment group showing enriched and depleted modules in vandetanib treated cells. (C) Gene set enrichment analysis comparing features of vandetanib enriched (resistant) module 1 and sensitive (depleted) module 6. (D) Single cell tamoxifen response score was higher in vandetanib resistant module supporting that dual treatment targets distinct cell populations. (E) Genes that were depleted in GDNF/RET high human tumors were depleted in vandetanib sensitive cells, supporting that signature as a biomarker of response. (****p<0.001; **p<0.01, Wilcoxon test.)
